# Biofilms promote tolerance and stability in gut bacterial communities during bile acid stress

**DOI:** 10.64898/2026.07.20.739518

**Authors:** Mónica Louro, Vitor Cabral, Karina B. Xavier

## Abstract

Several mechanisms have been described to explain how bacterial species colonize and persist in the mammalian gut. However, biofilm formation remains underexplored as a mechanism for gut microbiota symbiont persistence. While evidence of biofilm formation by individual gut symbionts is beginning to emerge, its occurrence and relevance in multispecies gut microbiota communities remain poorly studied. Here, we established an *in vitro* model to study multispecies gut microbiota biofilms using the Oligo-Mouse Microbiota 12 (OMM12) consortium, a defined community of murine gut isolates. We investigated community biofilm formation and responses to bile acids, host-derived detergent-like molecules released into the gut that can perturb bacterial growth and community structure. We identified distinct contributions of two OMM12 members: removal of *Enterococcus faecalis* strongly reduced community biofilm biomass, whereas removal of *Bacteroides caecimuris* had limited effect on biomass but strongly altered species associations. These results, together with monoculture assays, show that individual biofilm capacity does not directly predict community-level contribution. Moreover, planktonic and biofilm communities had markedly distinct responses to bile acid stress. Planktonic cultures, while more susceptible to bile acids, showed resilience by recovering growth yield within 24 hours upon bile stress removal. Community biofilms, in contrast, showed greater tolerance to bile acid stress and more species associations. Overall, our findings support biofilms as a community-level lifestyle that can buffer defined gut microbiota communities against host-associated chemical perturbations.

**Importance:** Despite decades of research, how the gut microbiota maintains diversity and persistence remains incompletely understood. Gut bacterial species must withstand harsh host-derived stresses while navigating complex interspecies interactions, many of which are highly competitive. In host-associated contexts, biofilms have largely been viewed as a detrimental trait to the host because of their role in pathogen persistence and protection from microbial clearance, leaving the potential contribution of commensal gut biofilms to microbiota stability underexplored. Our work establishes a simple and adaptable experimental framework for studying biofilm formation in a defined multispecies gut bacterial community. We show that biofilms alter how this community responds to bile acids, host-derived molecules that can disrupt bacterial growth and community structure. Our findings support biofilm formation as a protective lifestyle that can help gut symbionts withstand bile acid stress, raising the possibility that community biofilms contribute to microbiota persistence under chemical stress encountered in the host.

## INTRODUCTION

The mammalian gut microbiota is a complex host-associated microbial ecosystem in which thousands of bacteria and other microorganisms proliferate, cooperate, and compete for survival. The intricate interactions among these microorganisms shape the gut environment and, subsequently, affect host health (1–5). In terms of bacterial composition, the mammalian gut is mainly composed of Bacillota (formerly known as Firmicutes) and Bacteroidota (formerly known as Bacteroidetes) (6–9). These phyla, together with species from Pseudomonadota (formerly Proteobacteria), Actinomycetota (formerly Actinobacteria), and Verrucomicrobiota (formerly Verrucomicrobia), perform different functions. Such functions include fermentation of dietary nutrients, vitamin production, protection against pathogens, and development and stimulation of the immune system (3, 10, 11). To execute these functions, gut symbionts encode a diverse set of colonization mechanisms, including metabolic pathways, antimicrobial molecules, and adhesion factors, that help them to stably colonize the gut (12–16). A possible, yet underexplored, gut colonization factor is biofilm formation by gut microbiota symbionts (17–19). Biofilms are defined as multicellular structures of microorganisms in which cells are frequently embedded within a matrix of extracellular polymeric substances (EPS) and adhere to each other and/or to a surface (20, 21). Although commonly associated with disease and pathogenicity, biofilms are not exclusive to pathogens; in fact, most microbes found in nature exist in biofilms (20). The biofilm lifestyle confers several advantages to bacteria, including prolonged residence within a host or on abiotic surfaces, close association with other bacterial cells, and increased tolerance to external insults such as antibiotics and host immune responses (22–24). In other fields of research, such as industry or agriculture, biofilm formation is studied for its biotechnological potential. In these areas, biofilm-producing bacteria are sought for their resilience, capacity to colonize and persist, and their alternative metabolic products (25, 26). These properties suggest that biofilm formation by gut symbionts could promote resistance to perturbation and prolonged persistence within the host (18, 19, 27). A prevalent human gut symbiont, *Bacteroides thetaiotaomicron*, has recently been shown to adopt a biofilm lifestyle (28, 29). Interestingly, biofilm formation in this bacterium is induced by bile acids (29, 30), host-derived molecules secreted into the gut that contribute to digestion and lipid absorption, and can be modified by gut symbionts in different ways (31, 32). Given their detergent-like properties, bile acids can inhibit the growth of some bacteria; biofilm formation could be a way for gut symbionts to adapt to the presence of these molecules and other challenges in the gut environment (19). In fact, the role of bile in inducing biofilm formation has also been reported in other gut bacterial species (19, 33–36).

While evidence for biofilm formation by gut symbionts is beginning to emerge (19, 37–39), it remains unclear which members contribute to this lifestyle, and whether biofilms alter community responses to host-associated stresses such as bile acids (27). In this study, we use a defined community of 12 bacterial members derived from the murine gut, the Oligo-Mouse Microbiota 12 (OMM12) (40), as a model to study multispecies biofilms. This synthetic bacterial community, when originally assembled, was designed to provide a defined community for studying colonization resistance against *Salmonella* Typhimurium. This community includes one or more representatives of the five most abundant phyla in the mouse gut and has the advantage of all OMM12 members being culturable in laboratory single cultures, able to coexist in co-cultures, and capable of stably colonizing the mouse gut, with reproducible dynamics over time and even across different rodent facilities (40–42). Since its establishment, OMM12 has been used to study several aspects of gut microbiota function, including how diet, host factors, and interspecies interactions influence bacterial ecology and evolution (43), tumor formation (44), and immune-cell stimulation (45, 46). Recent studies have shown that OMM12 interspecies interactions are highly context-dependent, varying with nutrient availability, culture medium, and host environment (42, 47). *In vitro*, *Enterococcus faecalis* can shape community composition through metabolic interactions and enterocin-mediated inhibition, whereas *Bacteroides caecimuris* and *Blautia coccoides* can act as environment-dependent keystone species, particularly under polysaccharide-rich conditions. Importantly, these effects are not universal across conditions or gut regions, emphasizing that the contribution of each member depends on the ecological context (42, 47). However, these studies were performed mainly in planktonic cultures or in gnotobiotic mice, leaving unclear whether a structured biofilm lifestyle reshapes OMM12 members’ contributions, species associations, and tolerance to gut-relevant stress. Here, we addressed this gap by establishing an *in vitro* biofilm model for OMM12 and used it to compare member contributions, community structure, and bile acid responses between biofilm and planktonic lifestyles. Excluding two individual members revealed distinct roles: removing *E. faecalis* revealed its role as a major contributor to biofilm biomass in the OMM12 community, whereas removing *B. caecimuris* showed that its absence does not significantly affect biofilm biomass but is crucial for species associations within the biofilm context. Moreover, individual biofilm-forming capacity did not predict community-level contribution, suggesting that community context and interspecies associations shape OMM12 biofilm formation. Finally, we show that biofilm and planktonic communities respond differently to bile acid stress: planktonic cultures are sensitive but resilient, whereas biofilm communities are more tolerant, stable, and able to preserve or recover species associations. Together, our work identifies biofilms as a community-level lifestyle, which can promote tolerance of gut microbiota communities to environmental stresses, providing evidence for a possible role of multispecies biofilms as a mechanism contributing to gut microbiota persistence.

## RESULTS

### mGAM promotes higher biofilm biomass and community diversity than AAM medium

The OMM12 synthetic community (Fig.1A) has been used as a model to study the complex interactions among members of the gut microbiota. To better understand how this community behaves in a structured environment, we began by characterizing its ability to form biofilms and comparing it with free-living planktonic growth (Fig.1B; see OMM12 cocktail composition in Fig.S1A). We tested two media: Anaerobic *Akkermansia* medium (AAM), formulated by Derrien et al. (48) and used to establish the initial OMM12 community model (40), and modified Gifu Anaerobic Medium (mGAM), a rich medium commonly used for the growth of anaerobic gut bacteria (49). Our goal was to use a medium that enabled all community members to grow and coexist under biofilm conditions. Communities cultured under biofilm-promoting conditions were compared with communities cultured in shaking conditions for planktonic growth (Fig.1B). Biofilm biomass and planktonic growth yield was higher in mGAM than in AAM (Fig.1C), while qPCR quantification of total 16S ribosomal gene copy numbers, used here as a proxy for bacterial load, revealed no significant differences between cultures grown in either medium (Fig.1D). Interestingly, although all 12 members were detected in all conditions, biodiversity (measured by Inverse Simpson index, which considers species abundance and evenness) was significantly higher in biofilm cultures grown in mGAM in comparison to the other culture conditions, except planktonic cultures in AAM (Fig.1E). Principal Coordinate Analysis (PCoA) based on Bray-Curtis dissimilarity showed that planktonic and biofilm communities grouped separately in both media (Fig.1F). *Bacteroides caecimuris* (Bac) was the most abundant species overall in mGAM cultures (Fig.1G, left panel; see median values of absolute 16S DNA copy numbers of each member in Fig.S1B). *Enterococcus faecalis* (Enf) relative abundance was higher in biofilms than in planktonic cultures grown in mGAM (Fig.1G, left panel). However, its higher relative abundance in biofilms resulted from a decrease in the absolute abundance of the other community members, as *E. faecalis* absolute abundance was similar between biofilm and planktonic cultures (Fig.S1B). In AAM cultures, *E. faecalis* dominated the community in both lifestyles, followed by *Flavonifractor plautii* (Flp) (Fig.1G, right panel; absolute 16S DNA copy numbers in Fig.S1C). The low biodiversity obtained in AAM medium can be explained by the dominance of *E. faecalis*. Overall, although both media supported detectable growth of all members, as mGAM promoted greater biofilm biomass and higher biofilm community diversity than AAM, we chose to continue the study using mGAM as the culture medium.

**Fig. 1.**
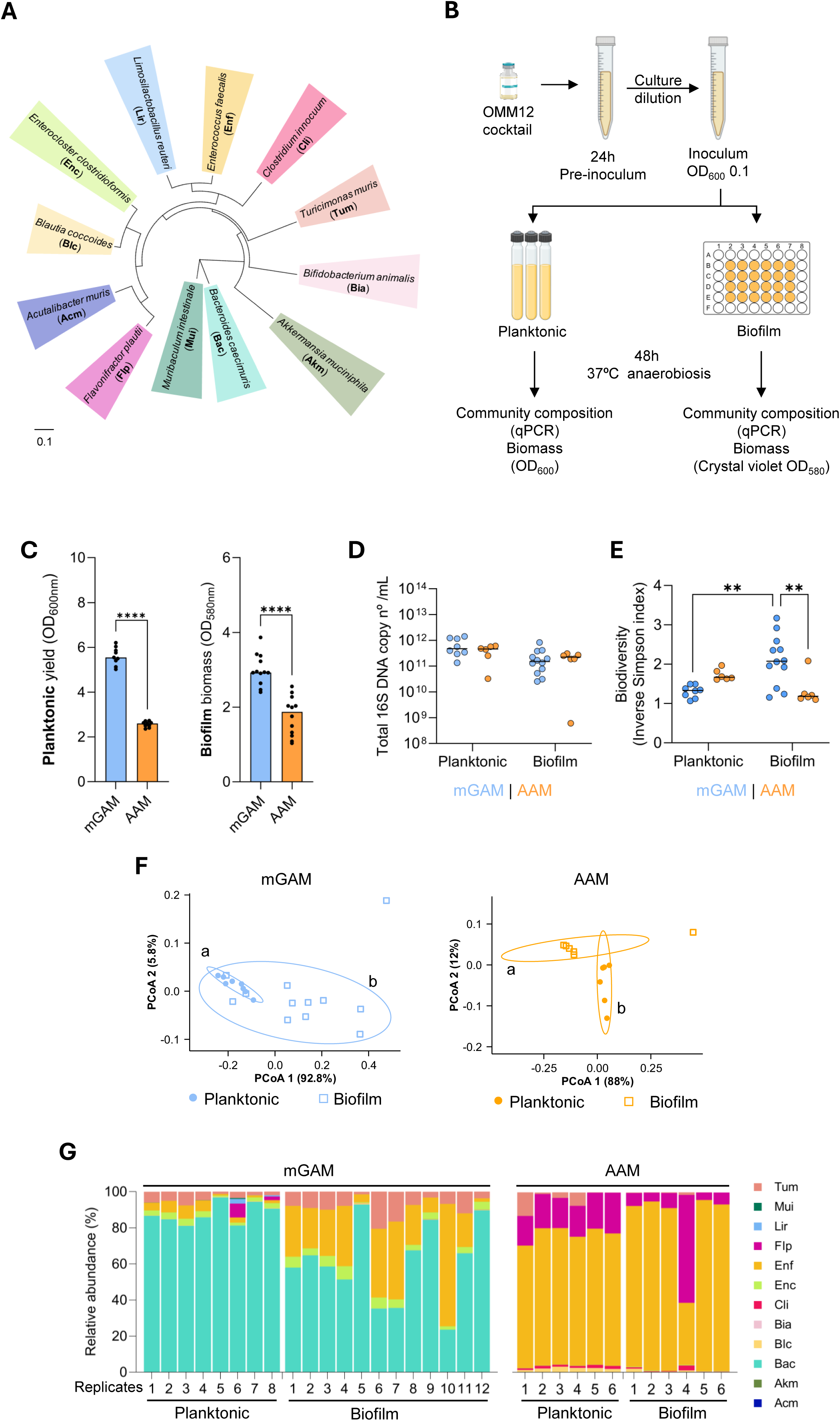
Biofilm biomass and diversity of OMM12 are higher in mGAM than in AAM. (**A**) Phylogenetic tree based on the 16S rRNA gene of species from the OMM12 synthetic community, and corresponding abbreviations and colour code. (**B**) Schematic overview of the experimental design. (**C**) OMM12 growth yield of planktonic cultures (OD600_nm_) and biofilm biomass (crystal violet assay (OD580_nm_) in mGAM and AAM, bars represent the median and dots represent the value of each replicate from 3 independent experiments, (**D**) concentration of total 16S DNA copy numbers, and (**E**) biodiversity (Inverse Simpson index) of cultures shown in **C**. (**F**) Principal Coordinate Analysis (PCoA) of Bray-Curtis dissimilarity index of planktonic and biofilm community relative abundance in mGAM and AAM, ellipses with a 90% confidence interval for the first two coordinates of each group were drawn on the associated PCoA. Each PCoA was tested with ANOSIM with Bonferroni correction. Different letters indicate significant differences between groups (p < 0.01). (**G**) Relative abundance of each species in planktonic and biofilm communities in mGAM and AAM, bars show the data for each culture replicate (total abundances are shown in **Fig. S1B** and **C**). In **C** data were analyzed with Mann-Whitney tests and **E** with Kruskal-Wallis’ test with Dunn’s correction for multiple comparisons (* p<0.05; ** p<0.01; **** p<0.0001).

### *Enterococcus faecalis* is a major contributor to biofilm formation in the OMM12 community

After establishing the experimental design and growth conditions for OMM12 biofilms, we sought to identify community members that contribute to biofilm formation. We first quantified biofilm formation by each OMM12 member in monocultures in mGAM. Among the 12 strains, *B. caecimuris* and *E. faecalis* produced the highest biofilm biomass (Fig. 2A). Because *B. caecimuris* was also the most abundant species in OMM12 cultures (Fig. S1B), and *E. faecalis* best discriminated the compositional shift between planktonic and biofilm cultures (Fig. S1D), we selected these two species for dropout experiments. We assembled two OMM11 communities: one without *B. caecimuris* (OMM-Bac; see cocktail composition in Fig.S2A) and another without *E. faecalis* (OMM-Enf; see cocktail composition in Fig.S2B). Excluding *B. caecimuris* had no significant effect on biofilm biomass, whereas removing *E. faecalis* caused a more than two-fold reduction compared with OMM12 (Fig.2B). Thus, although both species were strong biofilm formers in monoculture, only *E. faecalis* was required for full biofilm biomass in the community context.

**Fig. 2.**
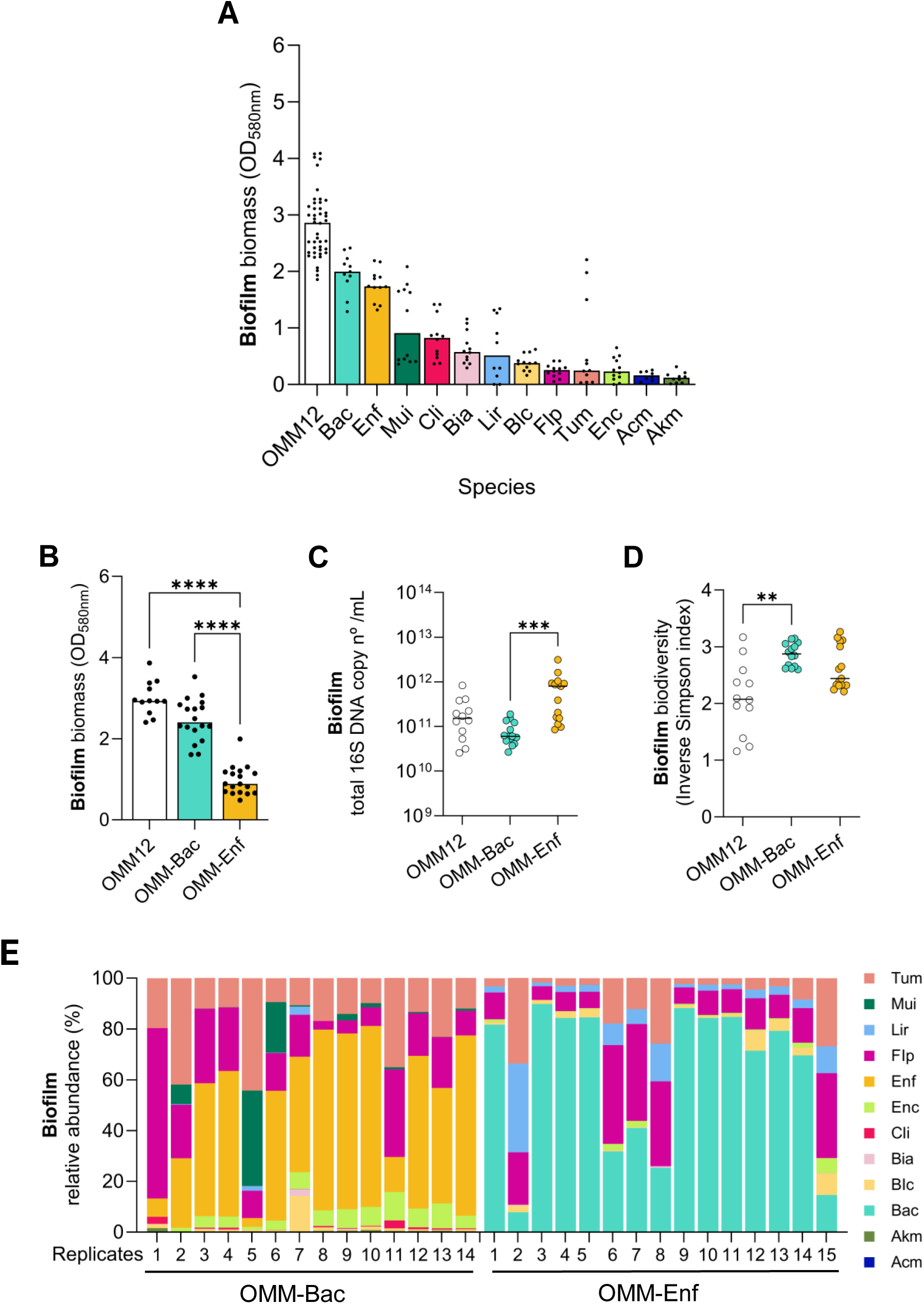
Removal of the two best biofilm formers *B. caecimuris* and *E. faecalis* reveals that absence of *E. faecalis* leads to decrease in biofilm community biomass without affecting ecosystem loads. (**A**) Biofilm biomass of OMM12 community and of each individual species in monocultures. (**B**) Biofilm biomass of OMM12 and OMM11 cultures, without *B. caecimuris* (OMM-Bac) or without *E. faecalis* (OMM-Enf). (**C**) Concentration of total 16S DNA copy numbers, and (**D**) biodiversity (Inverse Simpson index) of cultures shown in **B**. (**E**) Relative abundance of community members in OMM11 biofilms, bars show the data for each culture replicate (total abundances are shown in **Fig. S3B** and **D**). Bars in **A** and **B** represent the median, and dots represent the value of each replicate from 3 independent experiments. **A-D** data were analyzed with Kruskal-Wallis’ tests (* p<0.05; *** p<0.001; **** p<0.0001). The OMM12 data in **B**-**D** is from the experiment shown in **Fig1**.

Despite the reduction in crystal violet-stained biofilm biomass, total 16S DNA copy numbers in OMM-Enf biofilms were not significantly different from OMM12 biofilms (Fig. 2C). This indicates that removal of *E. faecalis* reduced biofilm-associated biomass without detectably affecting total bacterial loads (Fig.2C). In contrast, removal of *B. caecimuris* decreased total 16S DNA copy numbers in biofilm cultures and was associated with a more even contribution of the remaining species to community composition (Fig.2Cand 2E; Fig. S3B). Consistent with this shift in evenness, OMM-Bac biofilms had a higher biodiversity index than OMM12 biofilms (Fig.2D), suggesting that the dominance of *B. caecimuris* may have been limiting community evenness under these conditions. A similar increase in diversity was observed in planktonic OMM-Bac cultures (Fig.S2E).

Together, these results highlight the value of integrating phenotypic and compositional measurements. Removal of either *B. caecimuris* or *E. faecalis* altered community composition and species abundance (Fig.2E; Fig.S2F–H; Fig.S3), but these compositional changes did not always correspond to changes in biofilm biomass. In mGAM, *E. faecalis* emerged as a major contributor to OMM12 biofilm biomass, whereas *B. caecimuris* had a stronger effect on community load and evenness than on biomass itself.

### OMM biofilm communities are more tolerant to bile acids than OMM planktonic communities

For gut symbionts, being able to grow in the presence of bile acids is an important factor for gut colonization and can act as a source of stress, where some are sensitive while others are resistant. As biofilms are reported to be more tolerant to different stresses than planktonic cultures (50, 51), and bile can induce biofilm formation in some gut bacteria (19, 33, 34, 36, 52), we sought to determine how the OMM communities would respond to this host-related stress. To test this, we supplemented mGAM with 0.5% bile acids and allowed biofilm and planktonic cultures to grow. In planktonic conditions, all three communities had a significant decrease in growth yield when exposed to bile (Fig.3A). In contrast, biofilm cultures either maintained (OMM-Bac) or even increased (OMM12 and OMM-Enf) their biofilm biomass when exposed to bile (Fig.3B). These results show that biofilm formation by OMM12 and OMM11 communities was tolerant to, or enhanced by, bile acid stress, whereas planktonic cultures were affected by this stress. These changes in response to bile acids did not always translate into equivalent changes in the 16S DNA copy numbers. For planktonic cultures, 16S DNA copy numbers were not very affected by bile acids, except for the decrease observed in the OMM-Enf exposed to bile (Fig.3C). This indicates that in planktonic conditions, the reduction in culture yield upon bile exposure was not fully captured by total 16S DNA copy numbers, bile may have caused stress-induced changes in cell morphology or loss of cell viability, less reflected in the DNA copy numbers. Regarding biofilm cultures, for the OMM12 and OMM-Enf biofilms, exposure to bile led to a decrease in DNA copy numbers (Fig.3D). This reduction indicates that bile is affecting the loads of some species, particularly reflecting the decrease in *B. caecimuris* abundance across communities (Fig.S5–S6), while at the same time inducing biofilm formation, and thus maintaining or even enhancing biofilm biomass. This response is consistent with the prediction that biofilm cells are reacting to bile stress by inducing traits that produce biofilm biomass as a protective response. This response is not solely due to *E. faecalis,* as it occurred even in its absence; *i.e*., the OMM-Enf biofilm biomass was also higher in the presence of bile. It is also unlikely to be caused by *B. caecimuris,* as its monoculture did not increase in biomass when exposed to bile, and an OMM10 community without both *B. caecimuris* and *E. faecalis* also had more biomass in the presence of bile than in its absence (Fig.S7). Importantly, even though the biomass of the OMM-Enf biofilm was twofold higher with bile, it was still lower than the biomass of OMM12 and OMM-Bac biofilms. This result further supports the importance of *E. faecalis* in biofilm formation within the OMM12 community. In both OMM12 and OMM-Enf, *B. caecimuris* was the most affected species upon bile exposure, decreasing 100-fold in both lifestyles (Fig.S4; see absolute abundance in Fig.S5 and S6), which is in line with the growth inhibition observed on the *B. caecimuris* monocultures grown in mGAM with 0.5% bile (Fig.S8).

**Fig. 3.**
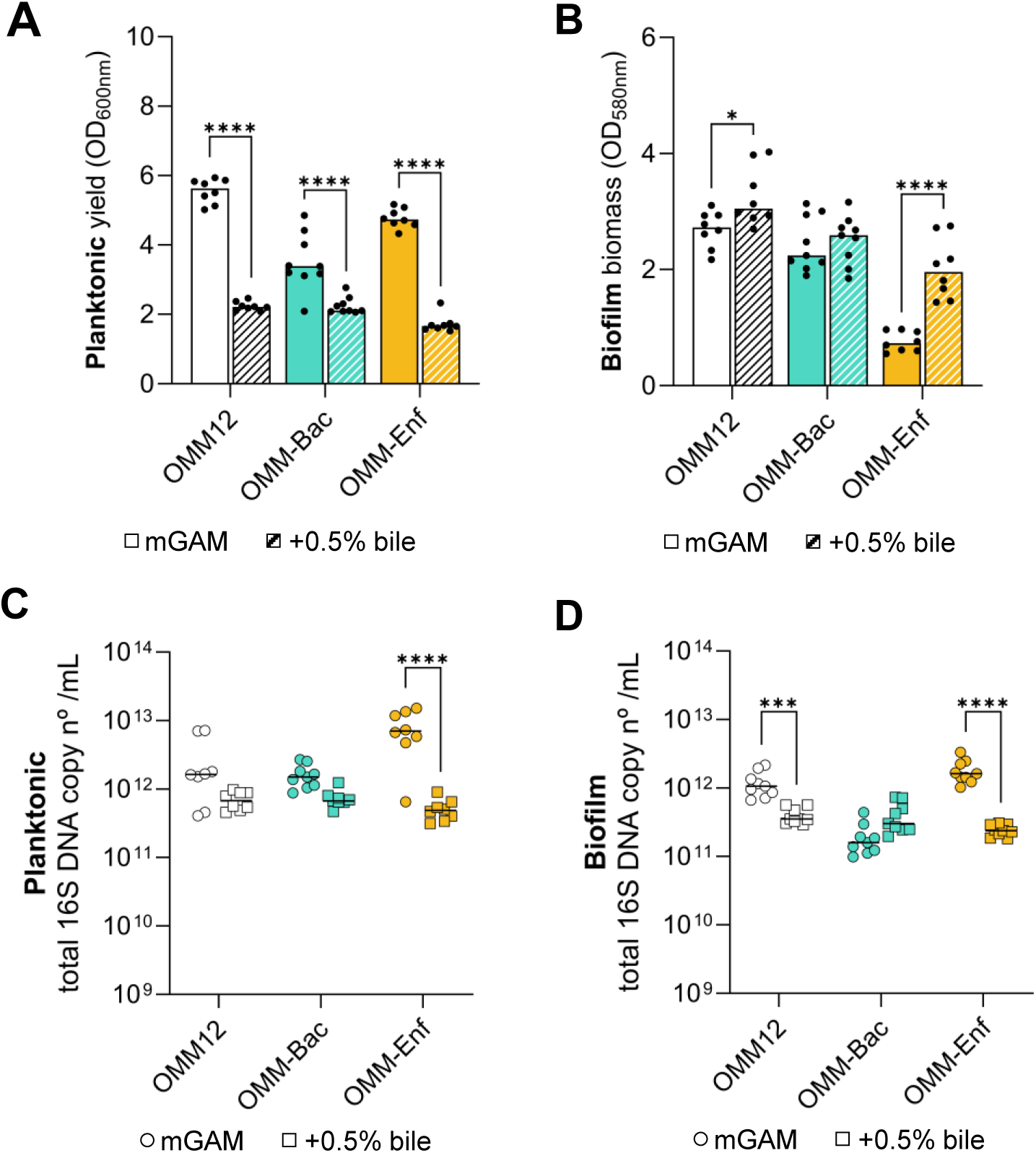
Biofilm communities are more tolerant to bile acid stress than planktonic communities. Effect of bile acids in OMM12 and OMM11 (**A**) planktonic and (**B**) biofilm cultures. Concentration of total 16S DNA copy numbers of (**C**) planktonic and (**D**) biofilm cultures in mGAM and mGAM with 0.5% bile of cultures shown in **A** and **B**, respectively. Bars in **A** and **B** represent the median, and dots represent the value of each replicate from 3 independent experiments. **A-D** data were analyzed with Mann-Whitney tests (* p<0.05; *** p<0.001; **** p<0.0001).

Altogether, these data show that biomass of biofilm cultures was tolerant to bile, whereas planktonic cultures’ growth yield was susceptible. In OMM12 and OMM-Enf communities, bile actually promoted biofilm biomass, with this effect being more pronounced in the OMM-Enf community. Moreover, OMM-Enf community had the lowest biofilm biomass in the presence of bile, again emphasizing the role of *E. faecalis* in biofilm formation by this community.

### Co-occurrence networks reveal more associations in OMM12 biofilms than in planktonic communities both in the absence and presence of bile

Having shown that planktonic and biofilm cultures respond differently to bile, we next sought evidence of potential differences in inferred associations among community members under biofilm and planktonic conditions, and asked how these associations changed under bile stress. We checked for co-occurrence patterns within the OMM12 and OMM11 planktonic and biofilm communities in the presence and absence of bile. Towards this goal, we used the absolute abundance data from the previous experiments to construct co-occurrence networks using Spearman correlations (Fig.4, Fig.S9 and 10). This method allowed us to infer possible positive, negative, or neutral associations – denoted as edges – between loads of the different species within these microbial communities. Since the communities do not have the same number of species – denoted as nodes – we normalized the number of edges by the number of nodes. OMM12 biofilms had the greatest number of inferred associations, consistent with the idea that a structured biofilm environment can promote stronger co-occurrence patterns among community members. Among the 12 conditions tested, OMM12 biofilms in mGAM had the highest ratio between number of edges and number of nodes (3.75, Fig.4). Strikingly, bile stress reduces these associations: OMM12 planktonic cultures exposed to bile lacked significant edges, but the reduction was smaller in biofilms (Fig.4 and Fig.S10A). Biofilms had 20 edges, 15 of which were the same as in the absence of bile (shared edges, Fig.S10B), thereby demonstrating biofilm robustness compared with planktonic cultures, similar to what was observed for biomass. We also observed species-dependent changes in network associations. The absence of either species (*B. caecimuris*, *E. faecalis*) in biofilm cultures resulted in fewer associations and lower edge-to-node ratios, in comparison to OMM12 (Fig.4). These results suggest that *B. caecimuris* contributes to the richness of inferred associations observed in OMM12 biofilms, as OMM-Bac biofilm communities had the lowest edge-to-node ratio (0.72, Fig.4) and shared fewer edges with OMM12 (Fig.S9C). *E. faecalis* appeared to have a smaller effect on overall network connectivity, although its absence was associated with novel positive co-occurrences in the absence of bile and novel negative co-occurrences upon bile stress (Fig.4 and Fig.S9-10). Moreover, we calculated the eigenvector centrality – a metric that weighs each connection by the importance of the neighbouring node; nodes connected to highly connected neighbours receive higher eigenvector scores, making this metric a sensitive indicator of potential network hubs in co-occurrence matrices (53). Eigenvector profiles of OMM12 and OMM-Bac biofilm communities support *B. caecimuris*’ role as a species promoting associations, with most species having scores above 0.8 in the case of OMM12 biofilm communities, while the OMM-Bac communities had only two or one species with scores above 0.8 in mGAM (Fig.S9D), in the absence or presence of bile (Fig.S10E), respectively. These results support the role of *B. caecimuris* as a putative biofilm-specific community network hub.

**Fig. 4.**
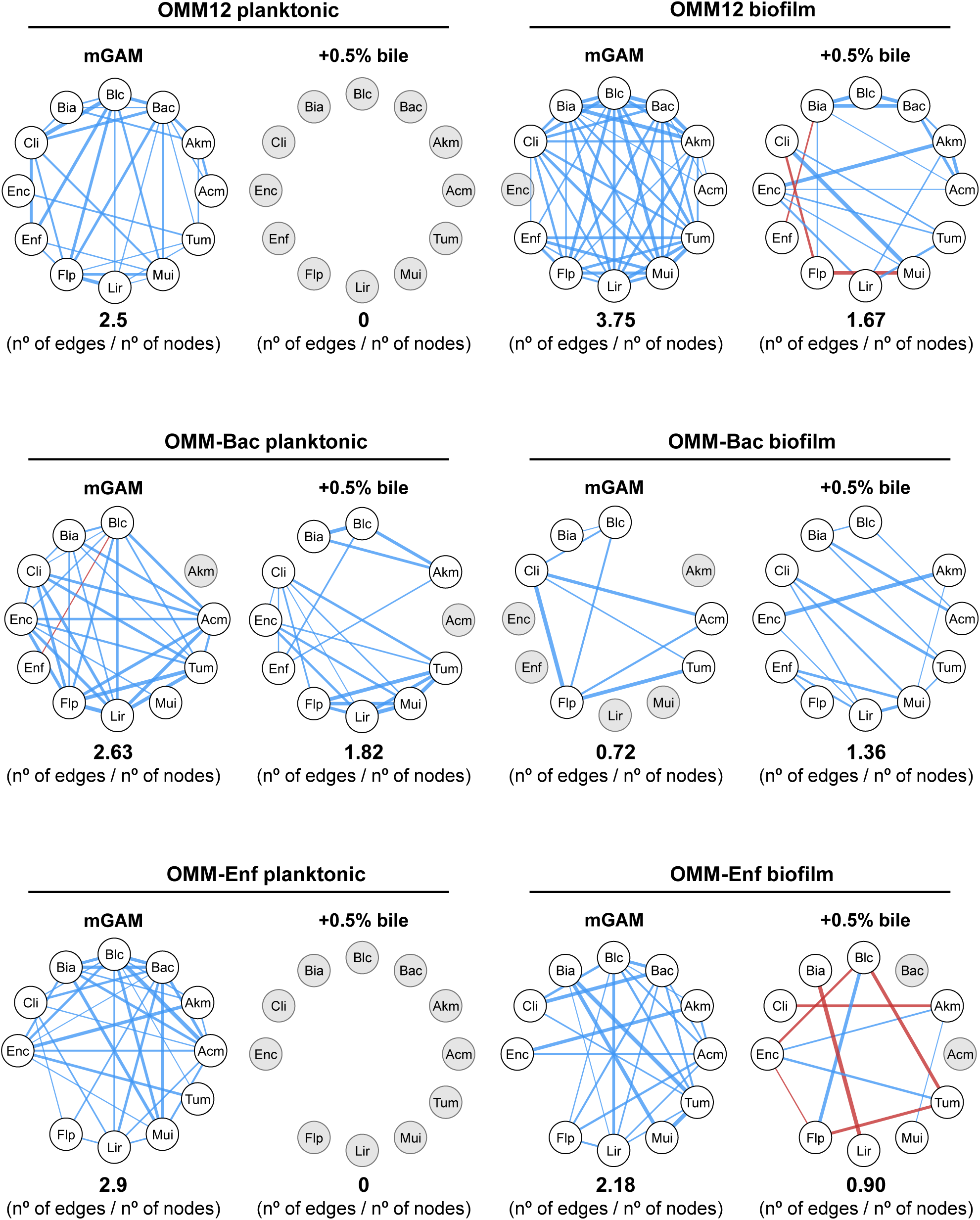
Co-occurrence network analysis in the presence and absence of bile identifies *B. caecimuris* as a network hub for species associations in OMM12 mGAM biofilms. Spearman correlation analysis network of OMM12, OMM-Bac and OMM-Enf planktonic and biofilm communities exposed to bile acids or not. Blue and red lines mean positive and negative correlation, respectively. Line thickness indicates strength of correlation. Significant edges are the ones whose |ρ| ≥ 0.6 and had a q-value ≤ 0.1. Grey nodes are nodes that did not have significant correlations with another member(s).

Taken together, these results indicate that OMM12 biofilms had more inferred interspecies co-occurrences and highly connected nodes than any other condition tested. Upon exposure to bile, OMM12 planktonic communities lost all significant associations, whereas OMM12 biofilm communities exposed to bile demonstrated greater robustness to the stress by keeping edges in common to mGAM, reflecting a greater robustness to stress. *B. caecimuris* had a high eigenvector centrality score, and its absence was associated with reduced network connectivity in mGAM biofilms, supporting its role as a putative community network hub under these conditions. Although *E. faecalis’* absence affected biofilm biomass, it did not affect the co-occurrence network to the same extent as *B. caecimuris* in mGAM conditions. In contrast, OMM-Enf communities were characterized by fewer positive associations and the emergence of novel negative connections. Collectively, while OMM12 biofilm results confirmed the expectation that biofilm communities are more tolerant to bile stress, the OMM-Bac and OMM-Enf results provide evidence that community context shapes interspecies associations in response to stress.

### Biofilms are tolerant and stable during bile stress, whereas planktonic cultures are bile-sensitive but resilient after stress removal

The effect of bile acids on community output was strongly lifestyle-dependent: biofilms were tolerant, maintaining or increasing crystal violet-stained biomass during bile exposure, whereas planktonic cultures were sensitive, showing reduced growth yield. Therefore, we next asked whether these communities could recover after bile acid stress was removed. After exposing communities to bile for 48 hours (48h stressed), the cell-free spent medium was replaced with fresh sterile mGAM with no bile and incubated for an additional 24 hours (+24h recovery). As a control, stressed cultures were compared with unstressed cultures grown in mGAM without bile for 48 h (48h unstressed), whose spent medium was also replaced with fresh mGAM and incubated for an additional 24 h (+24 h control). Community composition, and planktonic growth yield and biofilm biomass were assessed after the recovery or control period (+24h recovery/control) and compared with the corresponding stressed or unstressed cultures from the initial 48h period (Fig.5A).

**Fig. 5.**
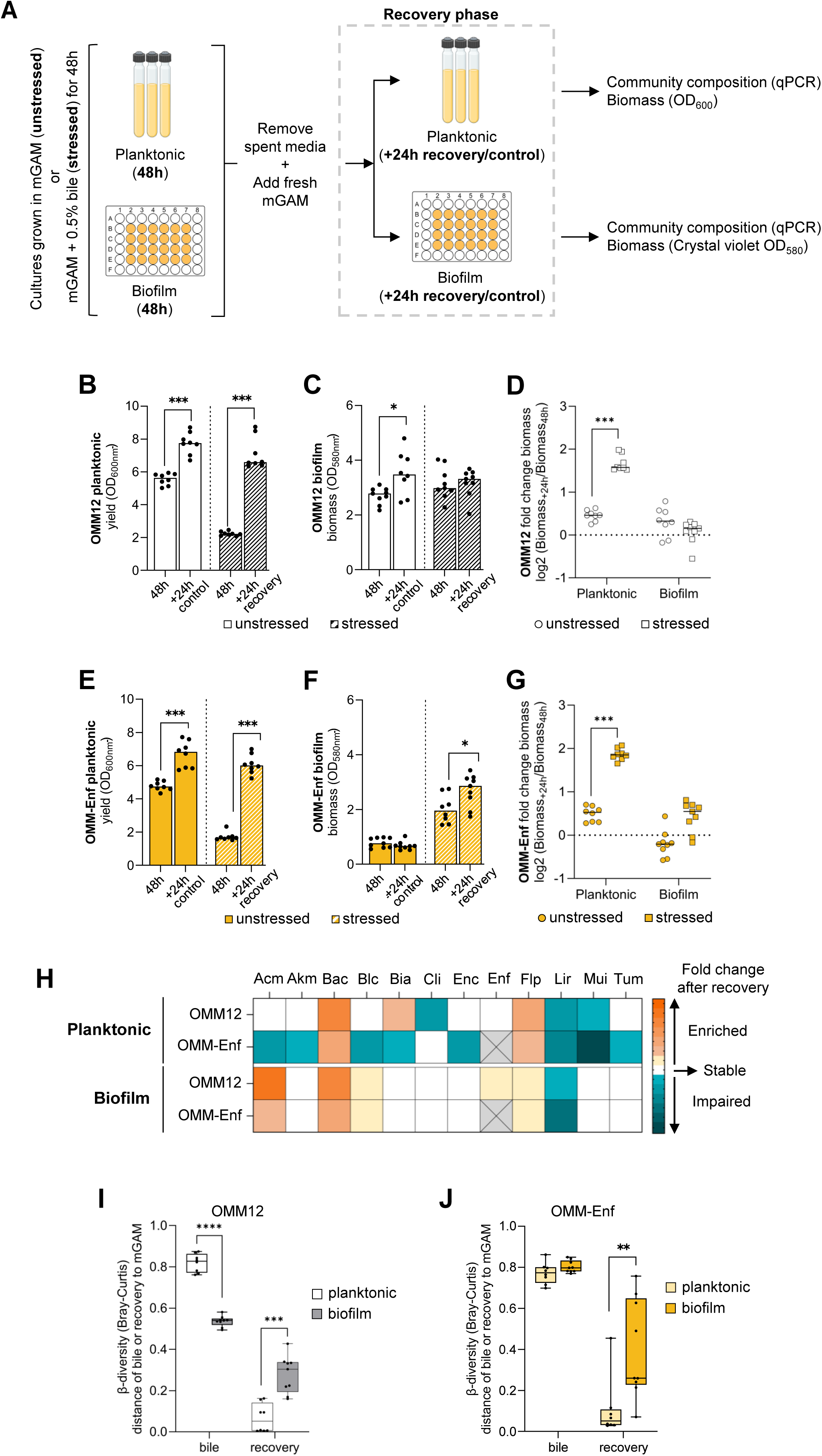
Biofilms are tolerant and stable to bile stress while planktonic cultures are resilient and fail to recover associations. (**A**) Schematic overview of the experimental design. (**B**) OMM12 planktonic growth yield and (**C**) biofilm biomass under unstressed (mGAM) and stressed (mGAM with 0.5% bile) growth conditions. (**D**) OMM12 fold change in planktonic growth yield and biofilm biomass cultures measured 24h after exposure to fresh media (unstressed) or after removal of bile acid exposure (stressed). (**E**) OMM-Enf planktonic growth yield and (**F**) biofilm biomass under unstressed (mGAM) and stressed (mGAM with 0.5% bile) growth conditions. (**G**) OMM-Enf fold change in planktonic growth yield and biofilm biomass cultures measured 24h after exposure to fresh media (unstressed) or after removal of bile acid exposure (stressed). (**H**) Heatmap representing the changes in abundance of each member after 24h recovering from bile acid stress; fold change above 1 was enriched, between -1 and 1 was stable and below -1 was impaired. (**I**) Analysis of the Bray-Curtis dissimilarity distances obtained comparing each lifestyle group to OMM12 mGAM 48h. (**J**) Analysis of the Bray-Curtis dissimilarity distances obtained comparing each lifestyle group to OMM-Enf mGAM 48h. Relevant PCoAs are shown in **Fig. S14**. Bars in **B**-**C** and **E-F** represent the median and dots represent the value of each replicate from 3 independent experiments; 48h data corresponds to the data shown in Fig.3A and **B**. In **B-G** and **I-J** data was analysed with Mann-Whitney tests (* p<0.05; ** p<0.01; *** p<0.001; **** p<0.0001).

We focused on the two communities with the most contrasting biofilm phenotypes: the complete OMM12 community, which formed high-biomass biofilms, and OMM-Enf, in which removal of *E. faecalis* impaired biofilm formation but bile acids still triggered a strong increase in biofilm biomass (Fig. 3B). Strikingly, OMM12 planktonic cultures showed resilience after stress removal, regaining growth yield within 24 hours to levels similar to unstressed controls (Fig.5B and D). This recovery in growth yield was accompanied by a slight, non-significant increase in total DNA copy numbers (Fig.S11A). In contrast, OMM12 biofilms were tolerant and stable, maintaining biofilm biomass and total 16S DNA copy numbers after bile removal (Fig.5C-D; Fig. S11B). Unstressed biofilm cultures, used as a control for comparison, showed a slight but significant increase in biomass production (Fig.5C-D) and in DNA copy numbers (Fig.S11B). Similar to OMM12, OMM-Enf planktonic cultures recovered growth yield after bile removal and maintained total 16S DNA copy numbers (Fig.5E and G; Fig. S11C). Regarding OMM-Enf biofilms, as with OMM12, biofilms showed greater tolerance than planktonic cultures, but in this case biofilm biomass continued to increase after bile removal (Fig. 5F-G; Fig. S11D).

We compared the abundance of each species after recovery (i.e., 24h after bile removal, with the abundance at the end of the bile treatment) and classified the log_2_ fold changes as follows: above 1 was considered enriched, between 1 and -1 stable, and below -1 impaired (Fig.5H). Regarding OMM12, biofilms had one impaired species, compared to three in planktonic (Fig.5H; see unstressed controls in Fig. S11E and community composition in Fig.S12A). Bray-Curtis dissimilarity index supported the overall stability of OMM12 biofilm community structure across bile exposure and recovery (Fig.5I; Fig.S13B and Fig.S14A-E). Under bile exposure, biofilm communities’ distance to the control was shorter than planktonic cultures’. By contrast, planktonic cultures shifted farther from the unstressed control during bile exposure but moved closer to the control than biofilms after bile removal, consistent with resilience rather than stress tolerance (Fig.5I; Fig.S12A and Fig.S13A). Although OMM-Enf planktonic cultures behaved similarly to OMM12 upon recovery in terms of growth yield (Fig.5E), community members were more severely impacted by bile acids, as 8 out of 11 species were impaired after recovery (Fig.5H). In contrast, OMM-Enf biofilm had more stable members (6 out of 11), and only one species remained impaired after stress removal (Fig.5H). Bray-Curtis distances again supported planktonic resilience: OMM-Enf planktonic cultures shifted away from the unstressed control during bile exposure but moved closer to it after recovery (Fig.5J and Fig.S12B, Fig.S13C and Fig.S14F-J). Compared with OMM12, OMM-Enf biofilm communities showed a more pronounced shift in community structure under bile stress, moving away from the unstressed control to an extent similar to that observed in planktonic cultures, and thus exhibiting weaker post-stress recovery, with high variation among replicates (Fig.5J). Together, these results indicate that removal of *E. faecalis* reduced biofilm community stability during and after bile acid stress. Concerning species associations, although planktonic cultures recovered growth yield and community composition in just 24h post-stress removal, we found no evidence that they recovered inferred species associations (Fig.6). In contrast, biofilms showed partial recovery of inferred species associations, with both OMM12 and OMM-Enf increasing their edge-to-node ratio and decreasing the number of negative edges after bile removal (Fig.6). Moreover, comparison of edge recovery suggested community-dependent restoration of inferred associations: OMM12 biofilms recovered 10 associations lost during bile treatment, whereas OMM-Enf recovered only 3 (Fig. 6; Fig.S15A,B). This further supports the idea that *E. faecalis* contributes to the stability and recovery of biofilm-associated species associations.

**Fig. 6.**
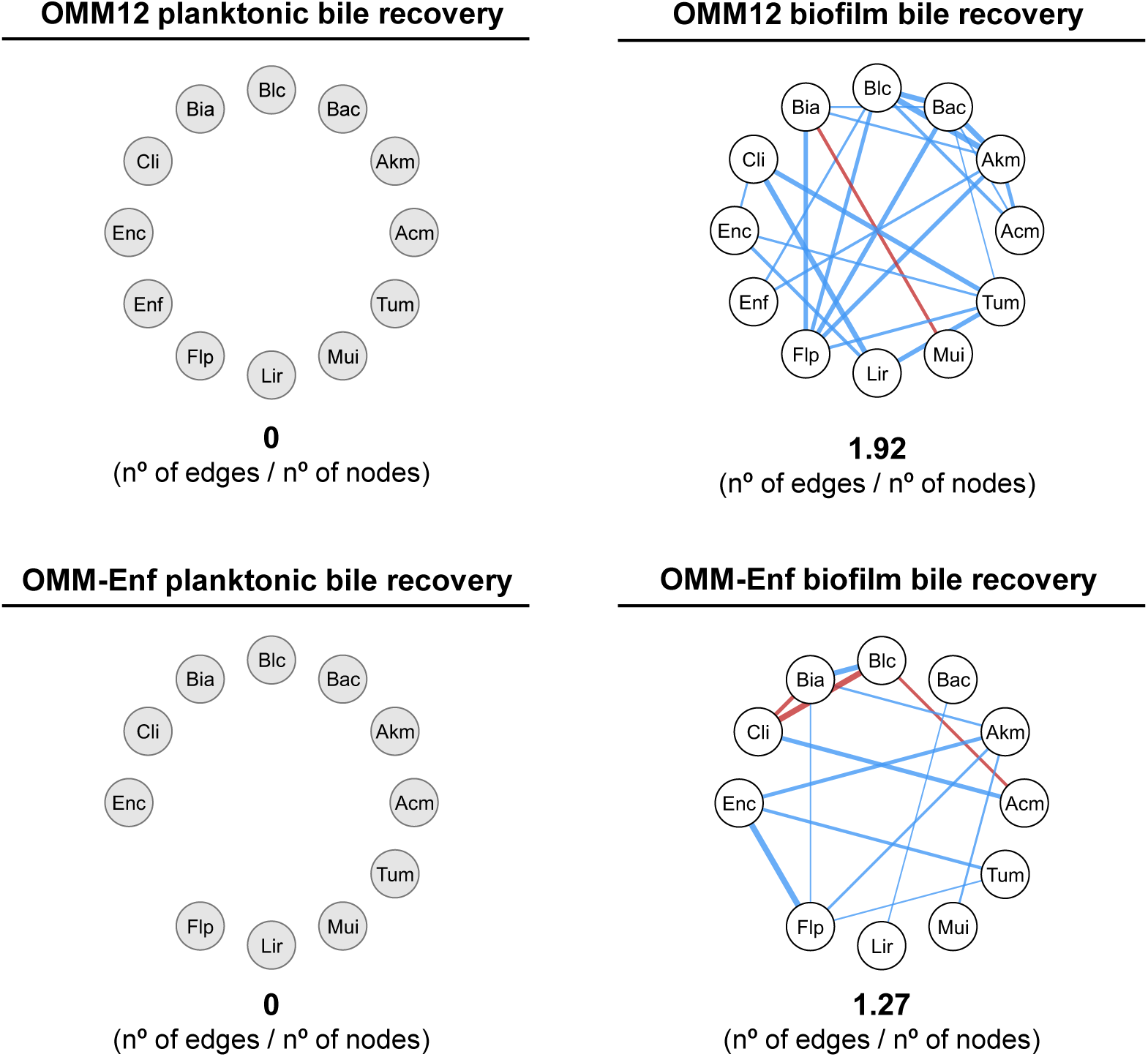
Biofilm cultures are more robust and show greater recovery of inferred associations lost during bile acid exposure than planktonic cultures. Spearman correlation analysis network of OMM12 and OMM-Enf planktonic and biofilm communities after bile acid removal. Blue and red lines mean positive and negative correlation, respectively. Line thickness indicates strength of correlation. Significant edges are the ones whose |ρ| ≥ 0.6 and had a q-value ≤ 0.1. Grey nodes are nodes that did not have significant correlations with another member(s).

In summary, planktonic cultures were sensitive to bile stress but resilient after stress removal, recovering growth yield and moving closer to unstressed community composition, while failing to recover inferred species associations. Overall, the presence of *E. faecalis* appeared to support biofilm community stability, as its absence reduced the recovery of species abundances. Both OMM12 and OMM-Enf biofilm communities were more tolerant to bile stress than their planktonic counterparts, maintained more stable species abundances after bile removal, and recovered some inferred associations lost during bile treatment. Taken together, these results show that the structured biofilm lifestyle promotes tolerance and stability during bile acid stress, whereas the planktonic lifestyle is more sensitive to stress but capable of rapid post-stress recovery.

## DISCUSSION

Biofilms can promote bacterial tolerance to environmental and host-associated challenges, but their relevance in multispecies gut microbiota communities remains poorly understood. Here, we investigated the ability of the OMM12 gut symbiont community to form multispecies biofilms and respond to bile acid stress. We found that OMM12 formed biofilms containing all 12 community members. *E. faecalis* was a major contributor to biofilm biomass, whereas *B. caecimuris* strongly affected community structure without affecting biofilm biomass. Consistent with a protective role for biofilms, biofilm biomass was maintained or increased during bile acid exposure in all communities tested, whereas planktonic cultures were bile-sensitive but resilient after stress removal.

Monoculture assays identified *B. caecimuris* and *E. faecalis* as the strongest individual biofilm formers, but their contributions in the community were markedly different. Exclusion of *E. faecalis* reduced OMM12 biofilm biomass without affecting planktonic yield, identifying this species as a major contributor to biofilm formation in the community context. In contrast, exclusion of *B. caecimuris* did not reduce biofilm biomass, despite its strong monoculture biofilm phenotype and high abundance in OMM12. Thus, as reported for other synthetic consortia (54–56), single-species biofilm behavior did not predict community-level contribution for the two strongest monoculture biofilm formers. These results suggest that OMM12 biofilm biomass emerges from community-level interactions that may include both positive and suppressive effects, with *E. faecalis* being necessary but not sufficient on its own for full community biofilm formation.

Our finding that *E. faecalis* is a major contributor to OMM12 biofilm biomass adds another layer to the context-dependent role of this species in defined gut communities. Weiss et al. showed that *E. faecalis* strongly affects OMM12 community composition *in vitro* under planktonic conditions, whereas its removal has a comparatively smaller effect in gnotobiotic mice, possibly because *E. faecalis* reaches low abundance under the *in vivo* conditions tested (42). At first glance, this could suggest that *E. faecalis*-dependent biofilm formation may have limited relevance for OMM12 colonization *in vivo*. However, our data indicate that the contribution of *E. faecalis* depends strongly on lifestyle: its removal had little effect on planktonic growth yield but markedly reduced biofilm biomass and altered biofilm-associated community behavior. This raises the possibility that the contribution of *E. faecalis in vivo* may become more apparent under host or dietary conditions that favor its expansion and increase exposure to stresses that can enhance biofilm-associated biomass. Because *E. faecalis* grows well on readily available carbohydrates such as simple sugars (42), Western-style diets enriched in simple sugars and fat (57) could favor its expansion. Moreover, because Western-style diets are associated with increased or altered bile acid pools (58, 59), and our data show that OMM-12-derived biofilm communities are tolerant to bile acids, it is tempting to speculate that *E. faecalis* and biofilm formation may contribute more strongly to community persistence under these dietary conditions. Although this remains to be tested *in vivo*, our results suggest that biofilm lifestyle, diet, and bile acid stress may interact to reveal context-specific roles of *E. faecalis* that are not evident under standard colonization conditions.

The distinction between biofilm biomass and community organization was further illustrated by *B. caecimuris*. Although its removal did not reduce total biofilm biomass, it caused several other members to increase in abundance and strongly reorganized inferred species associations within biofilms. This expansion of other community members in the absence of *B. caecimuris* suggests that, under mGAM conditions, *B. caecimuris* may act as a strong resource competitor, limiting the growth of species that rely on overlapping nutrients. However, a purely competitive role does not fully explain its position in the OMM12 biofilm network. OMM12 biofilms displayed the highest association-to-node ratio among all conditions tested, whereas OMM-Bac biofilms were among the least connected, suggesting that *B. caecimuris* also contributes to the structure of the inferred association network. One possible explanation is that *B. caecimuris* has a dual role: it may compete strongly for shared nutrients while also facilitating other members through metabolic by-products. Consistent with the carbohydrate-degrading capacity described for *B. caecimuris*, this species may release smaller oligo- and monosaccharides or fermentation products that can become available to other community members through cross-feeding (15, 60). In a structured biofilm, where cells remain in close proximity, such local by-product release could promote positive associations despite overall nutrient competition. This dual competitor–facilitator role could explain why *B. caecimuris* behaves as a putative network hub in OMM12 biofilms, while its removal markedly reduces inferred associations without reducing total biofilm biomass.

Bile acids shape gut microbiota composition, and gut bacteria must adapt to changes in host bile acid profiles. Because the OMM12 community encodes bile salt hydrolases but appears to lack pathways for converting primary into secondary bile acids (61), and the unconjugated primary acids it generates are toxic to many bacteria (62–64), we expected bile exposure to impose stress on this community. Consistent with this expectation, bile strongly reduced planktonic growth yield. In contrast, biofilm biomass was maintained or increased across all communities tested, supporting the idea that the biofilm lifestyle buffers OMM12-derived communities against bile acid stress. *E. faecalis* tolerance to bile may help explain why *E. faecalis*-containing communities maintained high biofilm biomass, but the increase in OMM-Enf biofilm biomass indicates that this response is not solely dependent on *E. faecalis*. The decrease in DNA copy numbers in OMM-Enf exposed to bile, together with the increased biofilm biomass, suggests that bile likely enhances matrix-associated biomass rather than bacterial load. This is consistent with previous reports that bile can induce biofilm formation in some gut bacteria (19, 33–36) and extends this observation to OMM12-derived multispecies communities.

Although biofilms buffered community-level biomass against bile stress, they did not protect all species equally. *B. caecimuris* was the member most strongly reduced by bile regardless of lifestyle, consistent with the reported sensitivity of *Bacteroides* species to unconjugated bile acids (62). This loss may have contributed to the disruption of inferred associations in OMM12 and OMM-Enf under bile stress, particularly if *B. caecimuris* contributes metabolites that support other members and their trophic interactions. By contrast, OMM-Bac retained more inferred associations under bile, suggesting that removal of *B. caecimuris* may relieve both competitive and facilitative dependencies and allow the remaining members to reorganize under stress.

After bile removal, the two lifestyles showed distinct recovery patterns. Planktonic cultures recovered growth yield and moved closer to unstressed community composition within 24 hours, but did not re-establish inferred species associations in OMM12 or OMM-Enf. Biofilms, in contrast, showed greater stability in species abundance across bile exposure and recovery, and recovered some associations lost during bile stress. In OMM-Enf biofilms, biomass continued to increase after stress removal, suggesting that bile exposure may initiate a response that persists after bile is removed. We therefore interpret biofilm communities as more tolerant and stable during bile acid stress, whereas planktonic communities are more susceptible during exposure but more resilient after stress removal.

Together, our results show that OMM12 biofilms represent a distinct community lifestyle in which member contributions, bile tolerance, stability, and inferred species associations differ from planktonic cultures. Importantly, species contributions could not be predicted from monoculture behavior or assumed to remain constant across community contexts. *E. faecalis* was a major contributor to biofilm biomass, whereas *B. caecimuris* shaped inferred association networks without being required for biomass. These findings highlight the value of studying gut bacterial communities in structured lifestyles and suggest that biofilm formation may contribute to microbiota persistence during host-associated chemical stress.

## MATERIALS AND METHODS

### Bacterial strains and media

The Oligo-Mouse-Microbiota 12 (OMM12) strains were acquired from the German Collection of Microorganisms and Cell Cultures (DSMZ). Each OMM12 member was cultured in 10 mL of selected medium (see below) to prepare frozen stocks. *Acutalibacter muris* (DSMZ 26090), *Akkermansia muciniphila* (DSMZ 26127), *Bifidobacterium animalis* spp. *animalis* (DSMZ 26074), *Clostridium innocuum* (DSMZ 26113), *Enterocloster clostridioformis* (DSMZ 26114), *Enterococcus faecalis* (DSMZ 32036), *Flavonifractor plautii* (DSMZ 26117), and *Turicimonas muris* (DSMZ 26109) were grown in a modified Anaerobic Akkermansia Medium (AAM; 18 gl^-1^ BHI, 15 gl^-1^ trypticase soy broth, 5 gl^-1^ yeast extract, 0.25 gl^−1^, 2.5 gl^−1^ K_2_HPO_4_, 1 mgl^−1^ haemin, 0.5 gl^−1^ D-glucose, 0.5 mgl^−1^ menadione, L-cysteine-HCl.H_2_O, 3% heat-inactivated fetal calf serum). *Bacteroides caecimuris* (DSMZ 26085), *Blautia coccoides* (DSMZ 29138), and *Muribaculum intestinale* (DSMZ 28989) were cultured in modified Gifu Anaerobic medium (mGAM, SHIMADZU). *Limosilactobacillus reuteri* (DSMZ 32035) was cultured in De Man-Rogosa-Sharpe medium (MRS, Gibco). Monocultures were incubated for 24 h, or 48 h in the case of *A. muris*, *A. muciniphila* and *T. muris*, at 37 °C without shaking under strictly anaerobic conditions (gas atmosphere 5% H_2_, 15% CO_2,_ and 80% N_2_). Monocultures were then subcultured in 1:10 dilution in 10 mL of mGAM medium and incubated in the same conditions for 24 h. Cultures were cryopreserved in glass vials with 1 volume of bacterial mixture to 1 volume of 40% glycerol solution supplemented with 0.2% L-cysteine-HCl · H_2_O and stored at -80 °C.

### Community assembly

To assemble the OMM12 community cocktail, the optical density at 600nm (OD_600_) of each monoculture was measured, and cultures were adjusted to 0.1 OD_600_, at a 1:1 ratio, except for *A. muris* and *T. muris*. These two species cannot be read at OD_600_ as they are translucent; instead, 2 mL of each was added to the cocktail. To assemble the OMM11 community cocktails, the protocol was the same, but no *B. caecimuris* or *E. faecalis* were added. Assembled communities were cryopreserved in glass vials with 1 volume of bacterial mixture to 1 volume of 40% glycerol solution supplemented with 0.2% L-cysteine-HCl · H_2_O and stored at -80 °C. An aliquot of the assemblies was used to extract DNA and assess community composition through qPCR.

### Biofilm cultures

Biofilm formation assays were performed by adapting a previously published method (65). Briefly, the bacterial cocktail was grown for 24 h in mGAM (or AAM where specified) at 37 °C without shaking under strictly anaerobic conditions. The following day, the culture was washed once in sterile PBS, and the final OD_600_ was adjusted to 0.1 in mGAM, and 500 μL of culture was dispensed per well in 48-well plates. Plates were centrifuged for 5 min at 4000 rpm at room temperature, and then incubated at 37 °C for 90 min under strict anaerobic conditions to promote adhesion. After this adhesion step, supernatants were carefully removed and discarded, wells were washed once with 500 μL of sterile PBS, and 1 mL of sterile fresh mGAM was carefully dispensed per well. Plates were incubated at 37 °C for 48 h under static and strictly anaerobic conditions. The attached biofilms (biomass) were quantified with Crystal Violet (CV). For that, supernatants were removed and discarded, and well bottoms were carefully washed with 500 µl of sterile PBS. Each well was stained with 0.1% CV solution, and plates were incubated at room temperature for 20 min, protected from light. After incubation, each well was washed with PBS, and plates were incubated open and inverted in a paper towel for 30 min, protected from light, to dry. Each well was then destained with 200 µl of a 33% glacial acetic acid solution and incubated for 15 min at room temperature, protected from light. Supernatants were removed from each well into a new plate and read for Abs_580_ in a Multiskan Sky plate reader (peak absorbance for CV staining), to quantify biofilm biomass. For the monospecies biofilm culture assays, the same protocol was followed.

### Planktonic cultures

For planktonic cultures, the bacterial cocktail was grown for 24 h in mGAM (or AAM, where specified) at 37 °C without shaking under strictly anaerobic conditions. The following day, the culture was washed once in sterile PBS, and the final OD_600_ was adjusted to 0.1 in mGAM, and 3 mL of culture was dispensed per Hungate tube. Tubes were incubated at 37 °C for 48 h with agitation (240 rpm) under strict anaerobic conditions. To quantify planktonic culture growth yield, 1 mL of culture was collected and centrifuged at 14000 rpm for 1 min. The supernatant was discarded, the pellet was washed with PBS and resuspended, and the optical density at 600 nm was measured.

### Bile acid stress and recovery assays

The different bacterial cocktails were grown for 24 h in mGAM at 37 °C, without shaking under strict anaerobic conditions. The following day, 1 mL of each culture was centrifuged at 6000 rpm for 90 sec, the supernatant discarded, and the pellet washed once in sterile PBS and resuspended. Then the optical density was measured, and a final OD_600_ was adjusted to 0.1 in mGAM or mGAM supplemented with 0.5% bile acids. One half of each culture was used for biofilm formation as described above, while the other half was used to grow the cultures in a planktonic lifestyle (with agitation at 240 rpm), for 48 h under strict anaerobic conditions at 37 °C. Biofilm and planktonic culture growth yield biomass were quantified as described above. For the recovery experiments, the supernatant from biofilms grown for 48 h in either mGAM or mGAM supplemented with 0.5% bile acids was collected from each well, washed once with sterile PBS, resuspended in fresh mGAM medium, and then carefully pipetted back into the original well. The plates were then incubated for another 24 h under strict anaerobic conditions. DNA samples were collected and crystal violet assays performed to assess biomass. Planktonic cultures grown for 48 h in either mGAM or mGAM supplemented with 0.5% bile acids were collected, washed once with sterile PBS, and resuspended in fresh mGAM medium. The tubes were then incubated for another 24 h under strict anaerobic conditions. DNA samples were extracted, and planktonic culture yield was assessed by OD600.

### Growth curve assays

All species, except *A. muris* and *T. muris*, for the reasons stated above, were grown in monoculture in mGAM for 24 h at 37 °C without shaking under strictly anaerobic conditions. The following day, the cultures were washed once in sterile PBS, and the final OD_600_ was adjusted to 0.1 in mGAM or mGAM supplemented with 0.5% bile acids. Then 150 µL of culture was dispensed into a 96-well plate. The plate was sealed to ensure an anaerobic environment and grown for 48 h without shaking, at 37 °C. Every 30 min, the plate was shaken and the OD600 measured using a Biotek Synergy H1 plate reader.

### DNA extraction and quantification

To assess community composition, 1 mL of sample was collected from planktonic and biofilm cultures (three to four wells of the biofilm cultures described above). In brief, DNA was extracted using the MasterPure Gram Positive DNA Purification kit (Lucigen/Biosearch technologies) following the manufacturer’s instructions to a final volume of 35 µL in DEPC water. DNA concentration was measured using QuBit BR DNA kit and adjusted to 1 ng/µL. For each sample, the 16S rRNA gene was amplified and quantified with species-specific primer pairs with the protocol adapted from B. Stecher’s group (40) under the following qPCR cycling conditions: 95 °C for 10 min, followed by 45 cycles of 95 °C for 15 s and 60 °C for 1 min for all strains. For *E. faecalis*, an annealing temperature of 65 °C was used to avoid nonspecific binding of DNA from other species. Standard curves were run for every qPCR measurement and were based on plasmids containing a single 16S copy for each of the 12 species. One PCR reaction (total volume of 8 µl) contained 0.16 µl of each primer at 10 μM, 0.032 µl of low ROX 50x, 4 µl of KAPA SYBR FAST mix buffer, 2µl of 1ng/mL gDNA and the remaining volume of DEPC water. qPCR reactions were run in duplicate.

### Co-occurrence network analysis

Absolute abundance data of each species were used to construct the co-occurrence networks. Pairwise co-occurrences between species were computed using Spearman’s rank correlation, which was selected for its robustness to non-normal distributions and outliers. From the adjacency matrix generated, only edges with |ρ| ≥ 0.6 were selected for further analysis. Network robustness was tested by performing 5000 permutations, which were then used to calculate an empirical p-value. P-values obtained were then adjusted for multiple testing using an FDR with Benjamini-Hochberg method threshold of 0.1. To check for outliers, a Leave-One-Out (LOO) analysis was performed, in which one replicate was left out and the analysis repeated iteratively for each replicate in the dataset. Inferred network hub nodes were defined as nodes with an eigenvector centrality score above 0.8. All analyses were performed in R (v4.4.3), using the package igraph (v2.1.4), and network graphs were constructed using the igraph and ggraph (v2.2.2) packages.

### Graphical representation and statistical analyses

All 16S DNA copy numbers obtained by qPCR were normalized to the expected number of 16S gene copies in each species’ genome and log_10_-transformed prior to plotting and statistical analysis. All graphs and respective statistical analyses were generated using GraphPad Prism 11 or R (v4.4.3), with the statistical tests specified in the figure legends and throughout the text. P values lower than 0.05 were considered significant and represented by asterisk symbols (\**p<0.05,* \*\**p*<0.01, \*\*\**p* <0.001, \*\*\*\**p* <0.0001). The Inverse Simpson diversity index was calculated using the R package vegan (v2.7-1). Bray-Curtis dissimilarity was computed from relative abundance data using the R packages vegan (v2.7-1) and ape (5.8-1). For Principal Component Analyses (PCoAs), statistical groupings were represented using a compact letter display, in which different letters indicate significant differences between groups (p < 0.05).

## Acknowledgments

We thank Raquel Gonçalo, Samuel Acheampong and Maria Ramirez-Montoya for helpful discussions, Ricardo Machado for reviewing the R scripts, Isabel Gordo, Mónica Serrano and Waldan Kwong for suggestions and helpful comments on the manuscript.

---

**Fig. S1.**
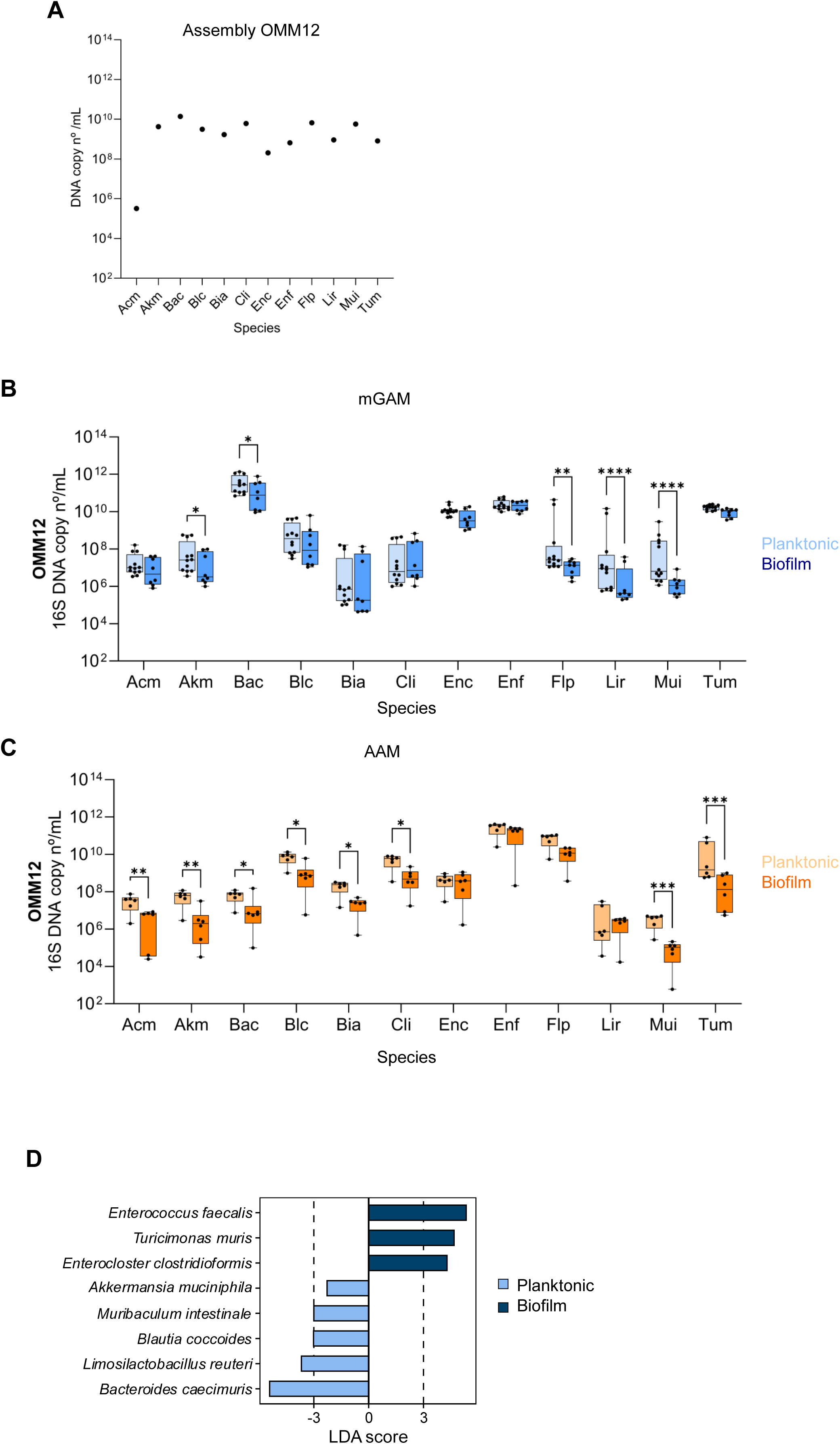
Absolute abundance of OMM12 communities. (**A**) Concentration of total 16S DNA copy numbers of assembled OMM12 cocktail. Absolute abundance of OMM12 biofilm and planktonic communities in (**B**) mGAM and (**C**) AAM, respectively, shown in Fig.1G. Dots represent the value of each replicate from 3 independent experiments. A linear mixed-effects model was used to determine statistical significance. Tukey adjustment was used to correct for multiple comparisons (* p<0.05; ** p<0.01; *** p<0.001; **** p<0.0001). Acm: *Acutalibacter muris*; Akm: *Akkermansia muciniphila*; Bac: *Bacteroides caecimuris*; Blc: *Blautia coccoides*; Bia: *Bifidobacterium animalis*; Cli: *Clostridium innocuum*; Enc: *Enterocloster clostridioformis*; Enf: *Enterococcus faecalis*; Flp: *Flavonifractor plautii*; Lir: *Limosilactobacillus reuteri*; Mui: *Muribaculum intestinale*; Tum: *Turicimonas muris*. (**D**) Linear discriminant analysis effect size (LEfSe) analysis identified species that differed significantly between planktonic and biofilm cultures. The horizontal bar chart shows LDA scores for species enriched in planktonic (light blue) and biofilm (dark blue). Only species with an LDA ≥ 2 are shown.

**Fig. S2.**
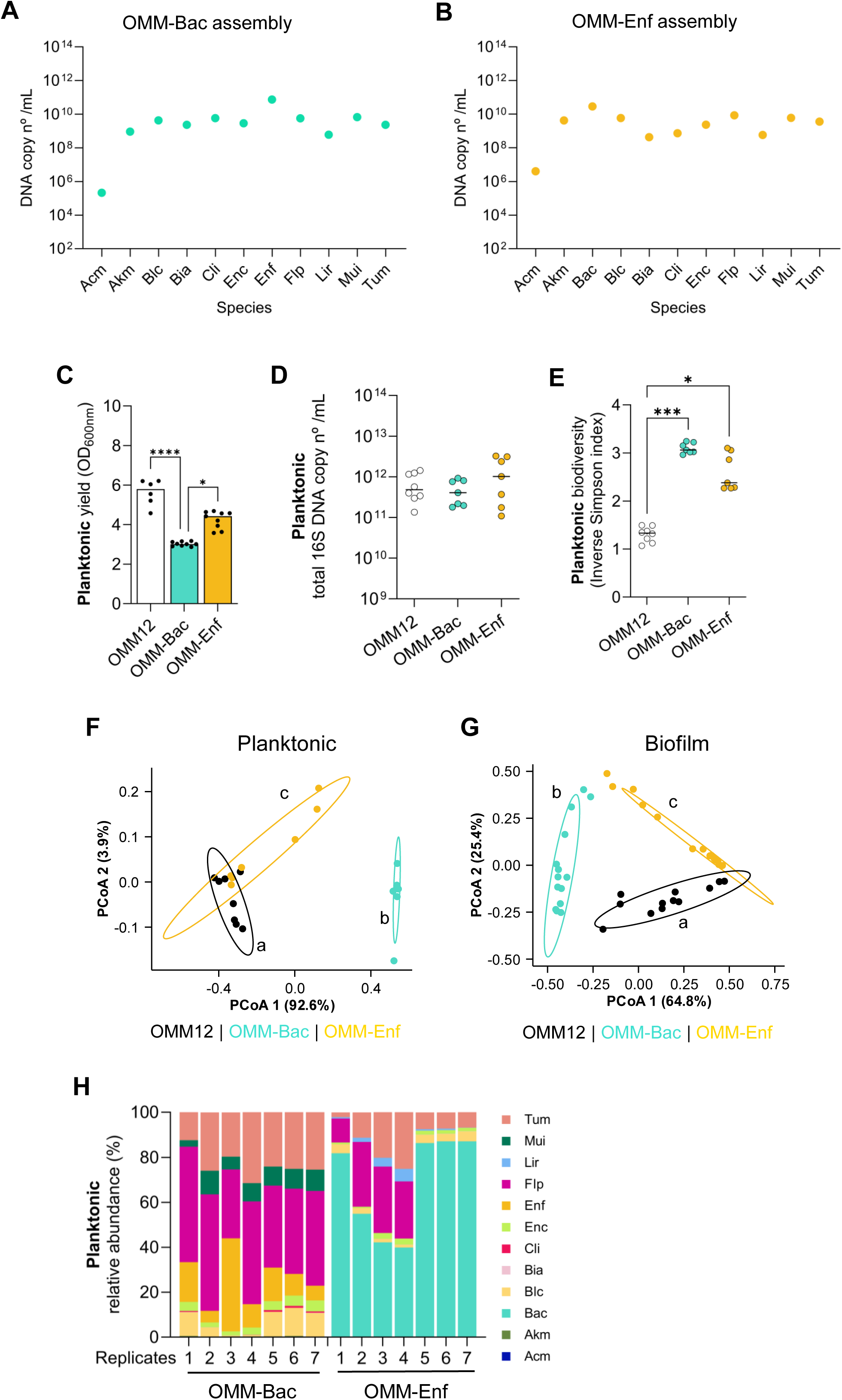
Planktonic data of OMM-Bac and OMM-Enf communities relative to Fig.2. (**A** and **B**) Concentration of total 16S DNA copy numbers of assembled OMM-Bac and OMM-Enf cocktails. (**C**) OMM112 and OMM11 communities planktonic cultures growth yield. (**D**) Concentration of total 16S DNA copy numbers and (**E**) biodiversity (Inverse Simpson index) of planktonic cultures. (**F**) Principal Coordinate Analysis (PCoA) of Bray-Curtis dissimilarity index of planktonic and (**G**) biofilm OMM12 and OMM11 communities; ellipses with a 90% confidence interval for the first two coordinates of each group were drawn on the associated plot. Each PCoA was tested with ANOSIM with Bonferroni correction. Different letters indicate significant differences between groups (p < 0.01). (**H**) Relative abundance of community members in OMM11 planktonic cultures, bars show the data for each culture replicate (total abundances are shown in **Fig.S3A** and **C**). **C-E** data were analyzed with Kruskal-Wallis tests (* p<0.05; *** p<0.001; **** p<0.0001).

**Fig. S3.**
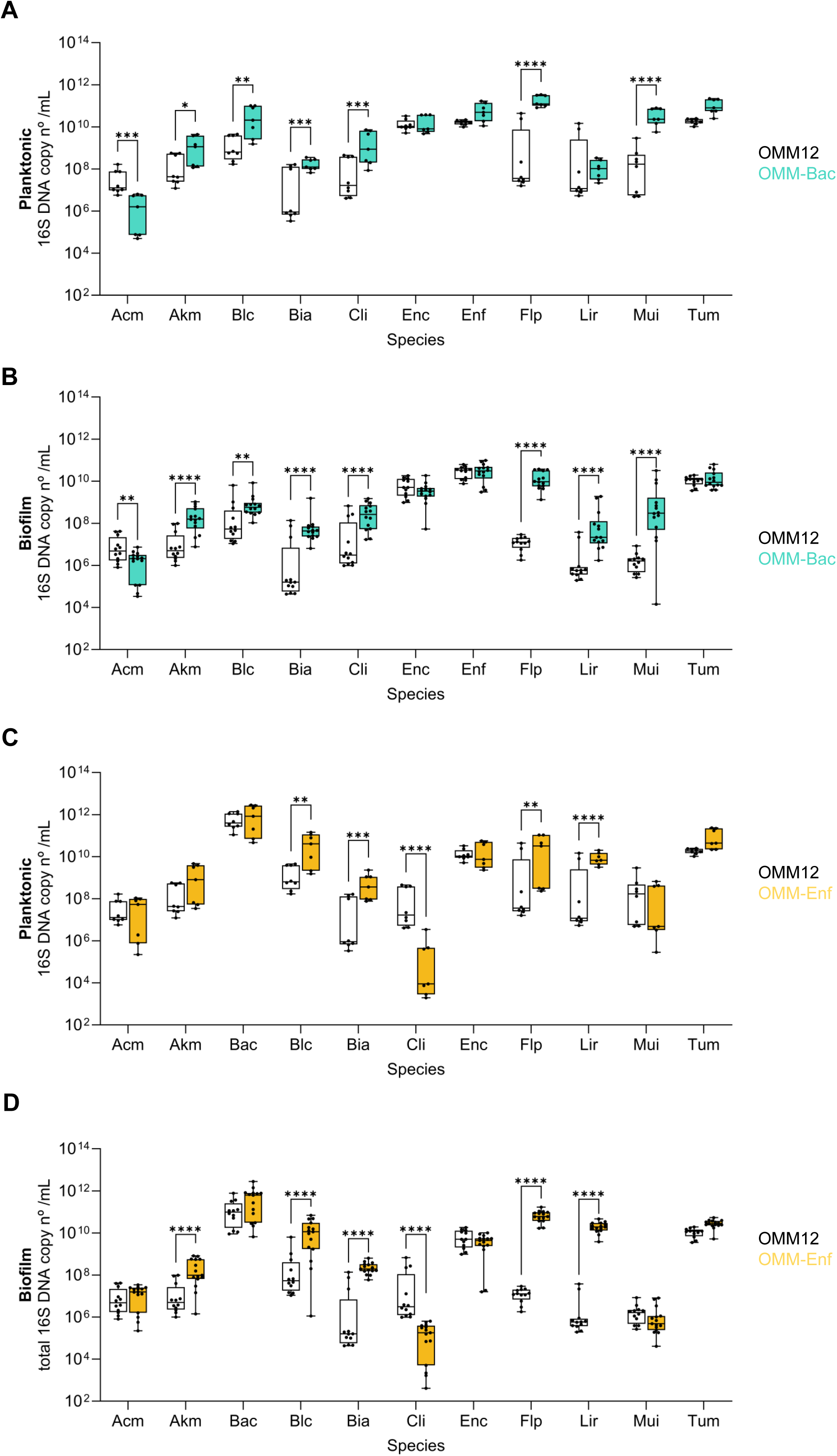
Absolute abundance of OMM-Bac and OMM-Enf communities shown in Fig.2B-E and Fig.S2. Absolute abundance of OMM-Bac (**A**) planktonic and (**B**) biofilm cultures shown in **Fig.S2** and Fig.2B**, 2E**, respectively, compared with absolute abundance of OMM12 cultures. Absolute abundance of (**C**) OMM-Enf planktonic and (**D**) biofilm cultures shown in **Fig.S2** and Fig.2B**, 2E**, respectively, compared with absolute abundance of OMM12 cultures. Dots represent the value of each replicate from 3 independent experiments. Comparison between each members’ abundance in OMM12 and OMM11 communities was analyzed with a linear mixed-effects model to determine statistical significance. Tukey adjustment was used to correct for multiple comparisons (* p<0.05; ** p<0.01; *** p<0.001; **** p<0.0001).

**Fig. S4.**
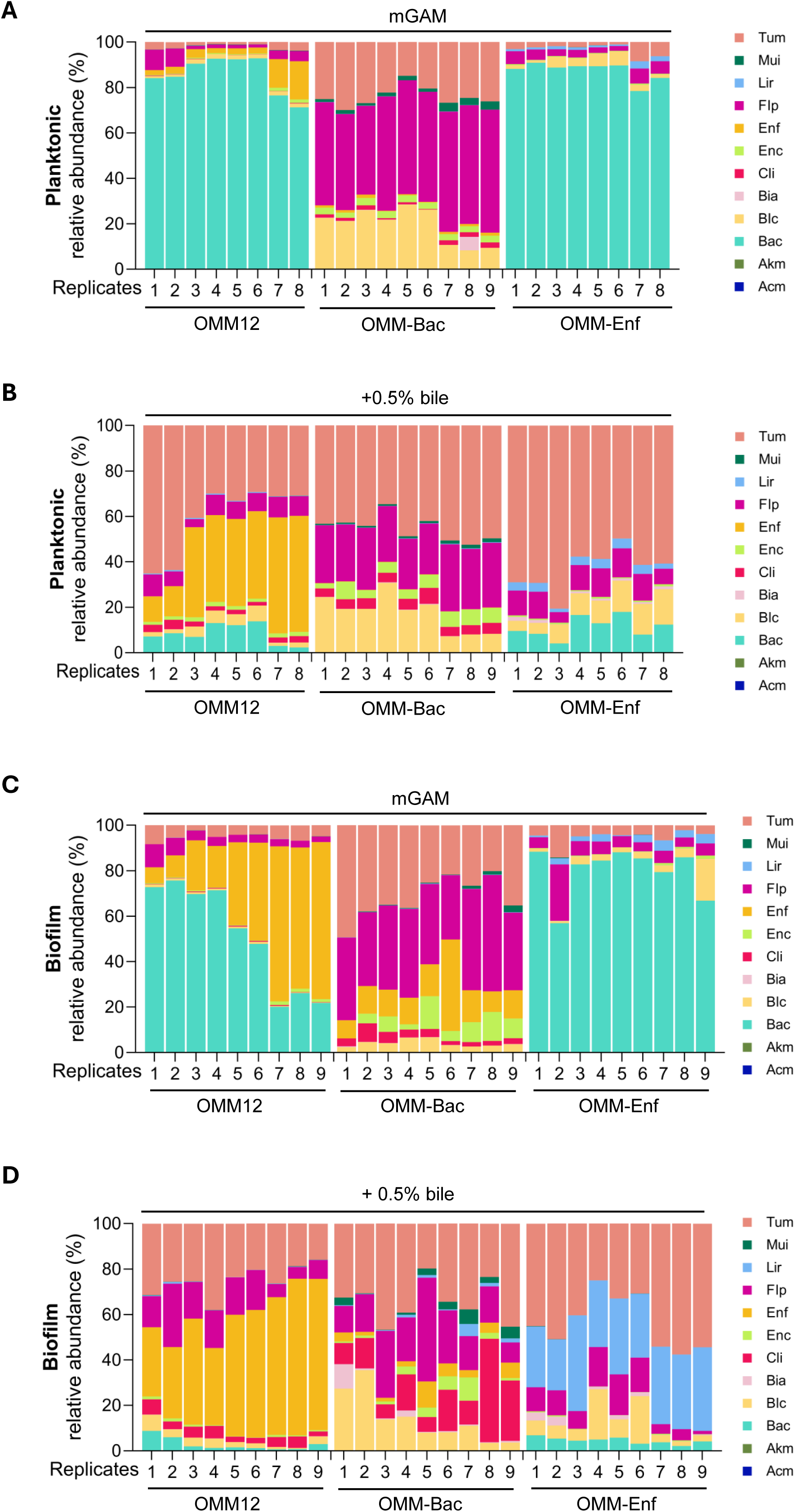
Relative abundance of OMM12 and OMM11 communities from in Fig.3. Relative abundance of planktonic (**A** and **B**) and biofilm (**C** and **D**) communities in mGAM and mGAM with 0.5% bile, respectively. Bars show the data for each culture replicate (total abundances are shown in **Fig.S5** and **6**).

**Fig. S5.**
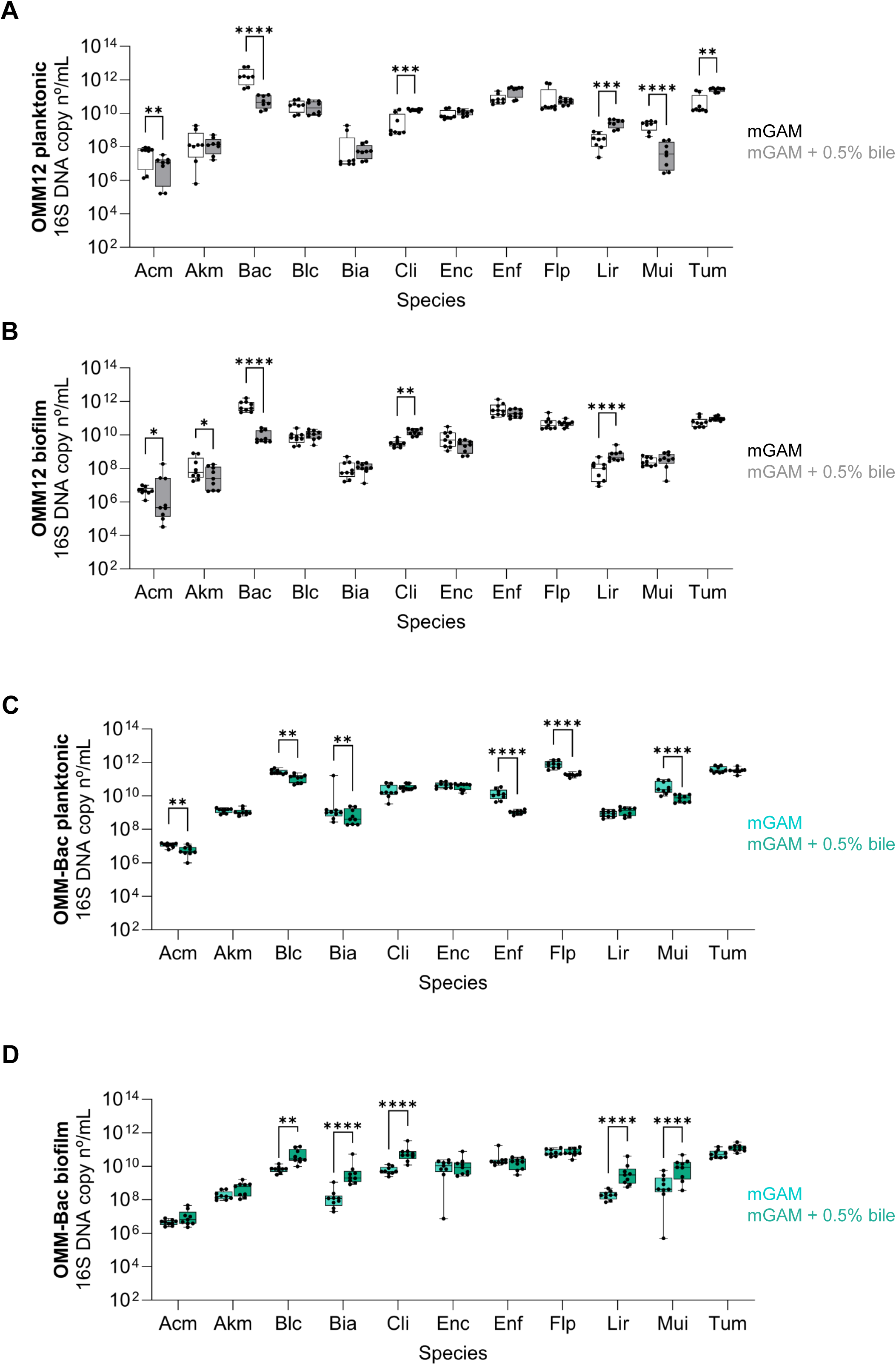
Absolute abundance of OMM12 and OMM-Bac communities shown in Fig.3. (**A**) OMM12 absolute abundance of planktonic and (**B**) biofilm communities. (**C**) OMM-Bac absolute abundance of planktonic and (**D**) biofilm communities. Each dot represents the data from a culture replicate from 3 independent experiments. Comparison between each member’s abundance in mGAM and bile exposed communities was analyzed with a linear mixed-effects model to determine statistical significance. Tukey adjustment was used to correct for multiple comparisons (* p<0.05; ** p<0.01; *** p<0.001; **** p<0.0001).

**Fig. S6.**
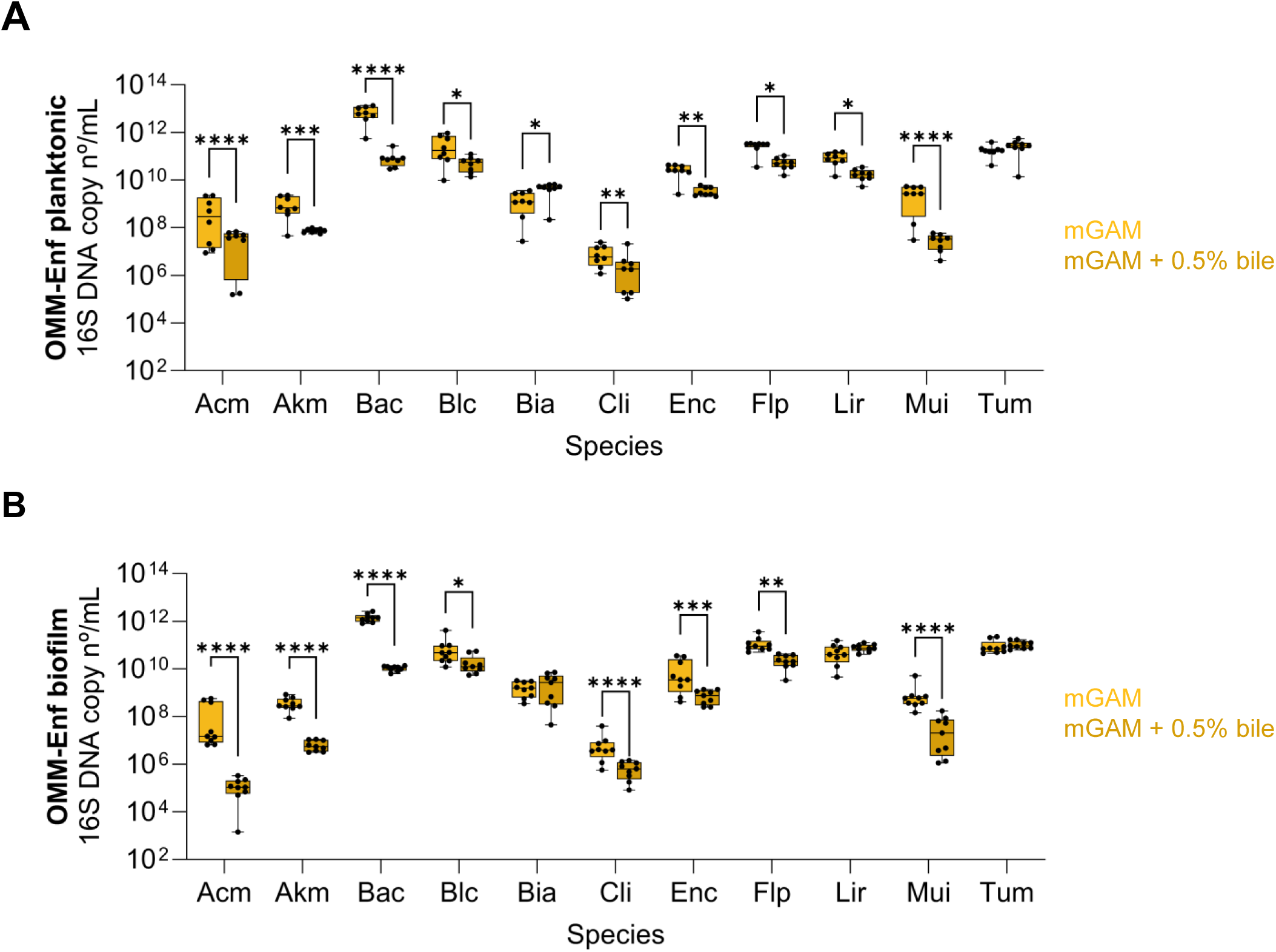
Absolute abundance of OMM-Enf communities shown in Fig.4. (**A**) OMM-Enf absolute abundance of planktonic and biofilm (**B**) communities. Each dot represents the data from a culture replicate from 3 independent experiments. Comparison between each member’s abundance in mGAM and bile exposed communities was analyzed with a linear mixed-effects model to determine statistical significance. Tukey adjustment was used to correct for multiple comparisons (* p<0.05; ** p<0.01; *** p<0.001; **** p<0.0001).

**Fig. S7.**
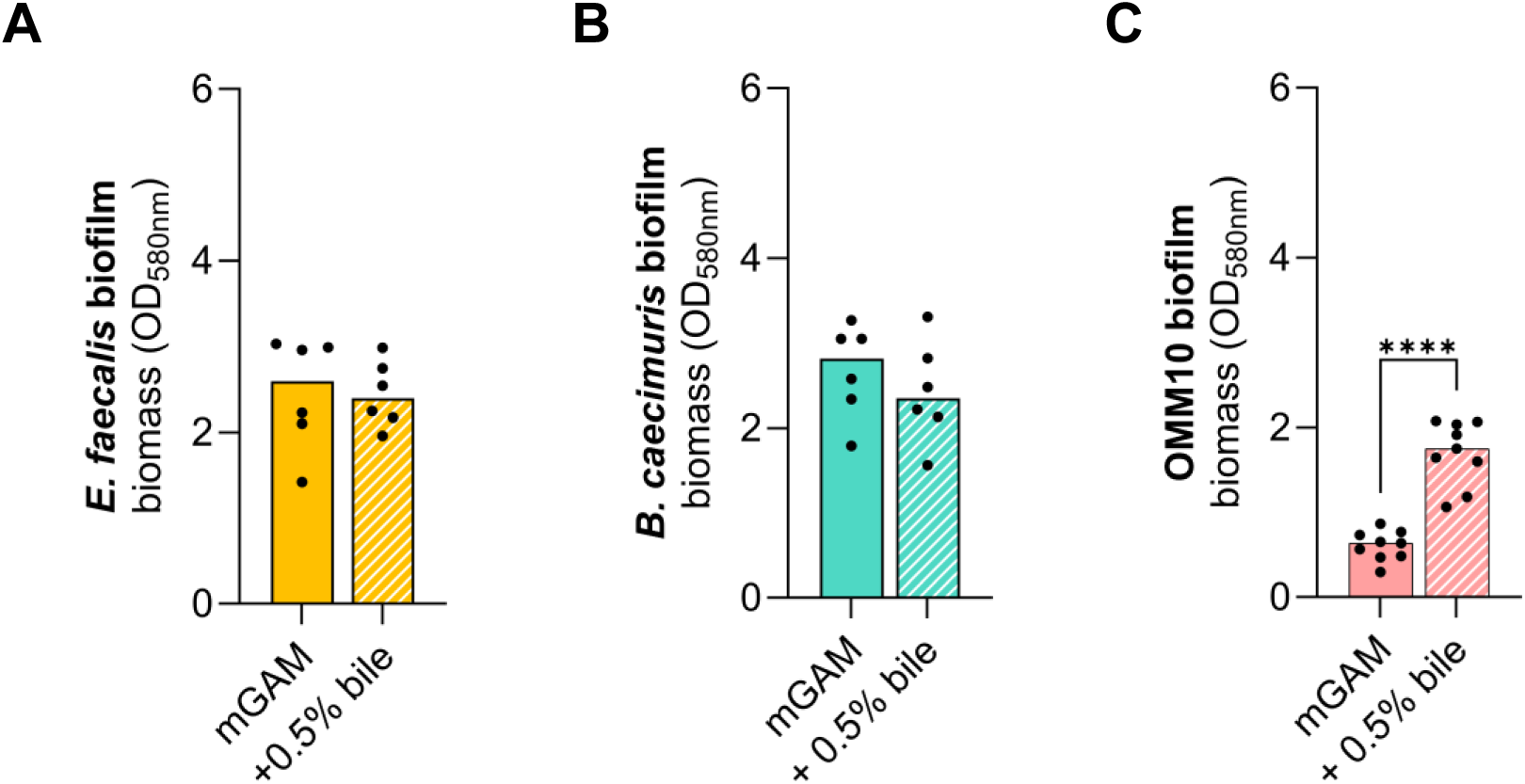
Bile does not affect *E. faecalis* and *B. caecimuris* biofilm monocultures. (**A**) Biofilm biomass of *E. faecalis* and (**B**) *B. caecimuris* monocultures and (**C**) OMM10 community exposed to bile acids. Data were analyzed with Mann-Whitney test (**** p<0.0001).

**Fig. S8.**
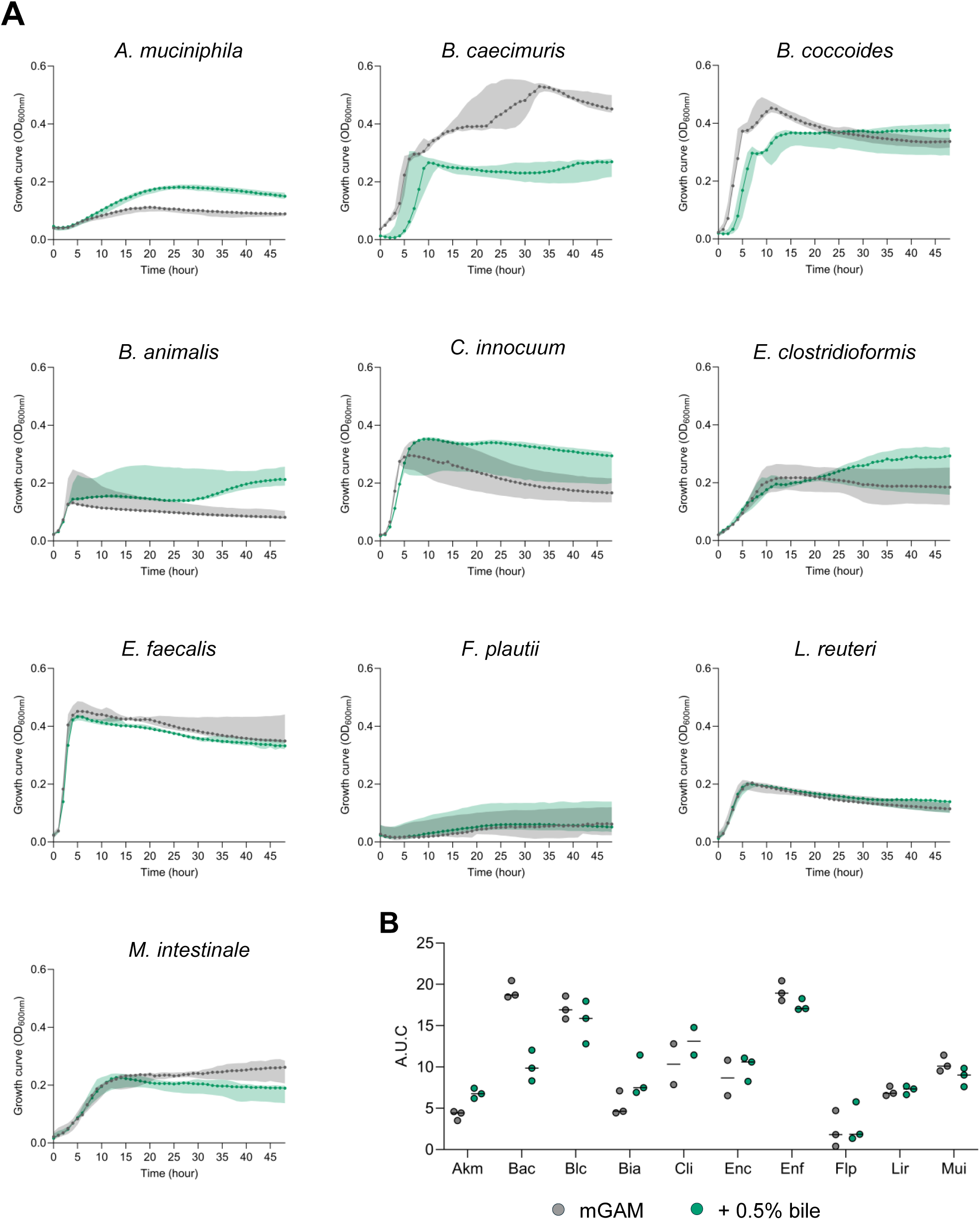
Monoculture growth with or without bile supplementation in mGAM. (**A**) 48h monoculture growth curves in mGAM (black) or mGAM with 0.5% bile (green). Shaded areas correspond to the standard deviation. (**B**) Area under the curve (A.U.C) of monoculture growth curves in **A**. Each dot corresponds to 3 independent experiments, from which 5 to 6 data points were collected.

**Fig. S9.**
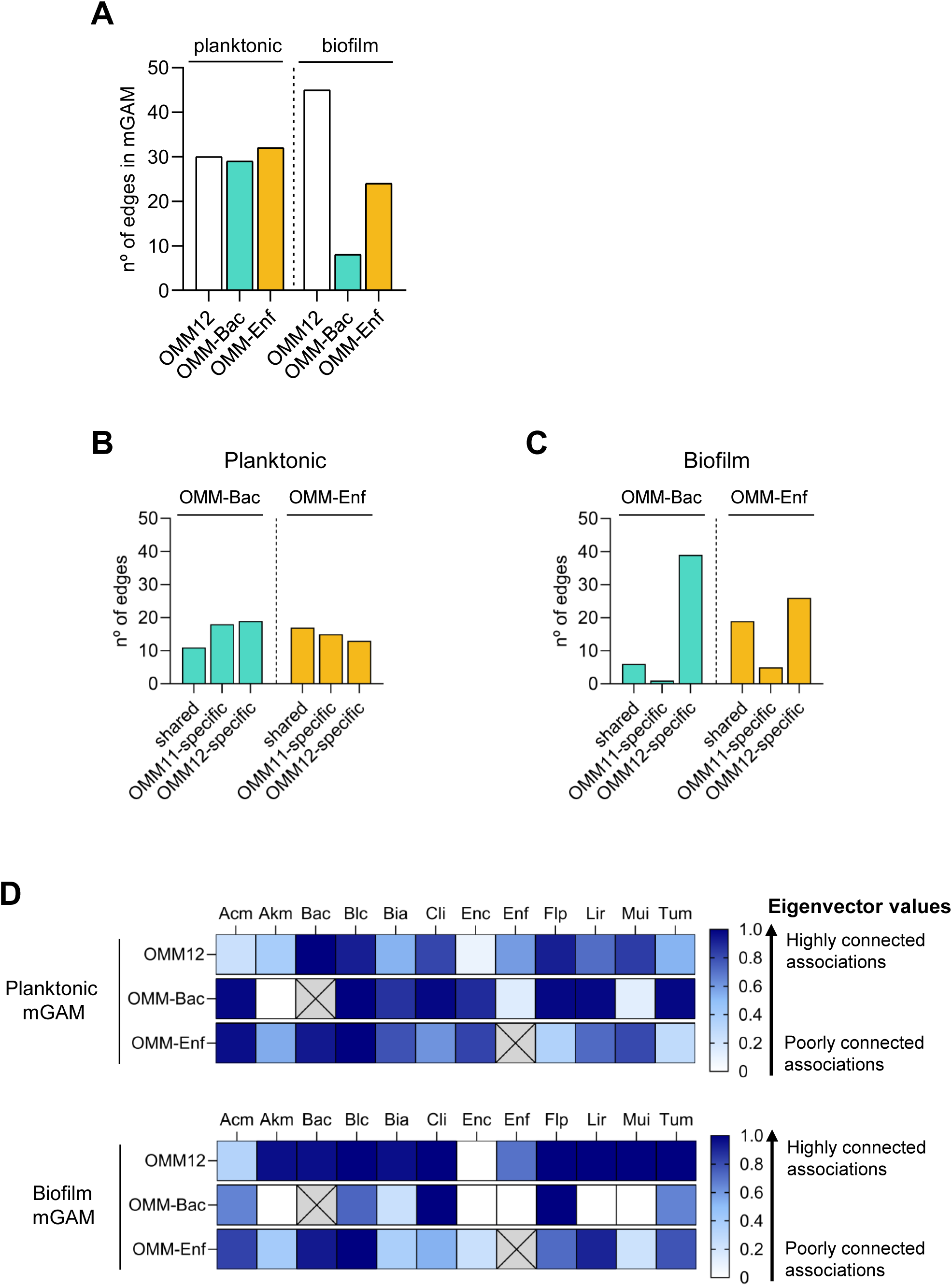
Network analysis of OMM12 and OMM11 communities shown in Fig.4 in mGAM conditions. (**A**) Total number of edges in OMM12, OMM-Bac and OMM-Enf planktonic and biofilm communities. (**B**) Comparison of number of edge pairs shared between OMM12 and OMM11 communities in planktonic and (**C**) biofilm. (**D**) Heatmap of eigenvector values that describe node centrality of each species of OMM12 and OMM11 planktonic and biofilm communities in mGAM conditions.

**Fig. S10.**
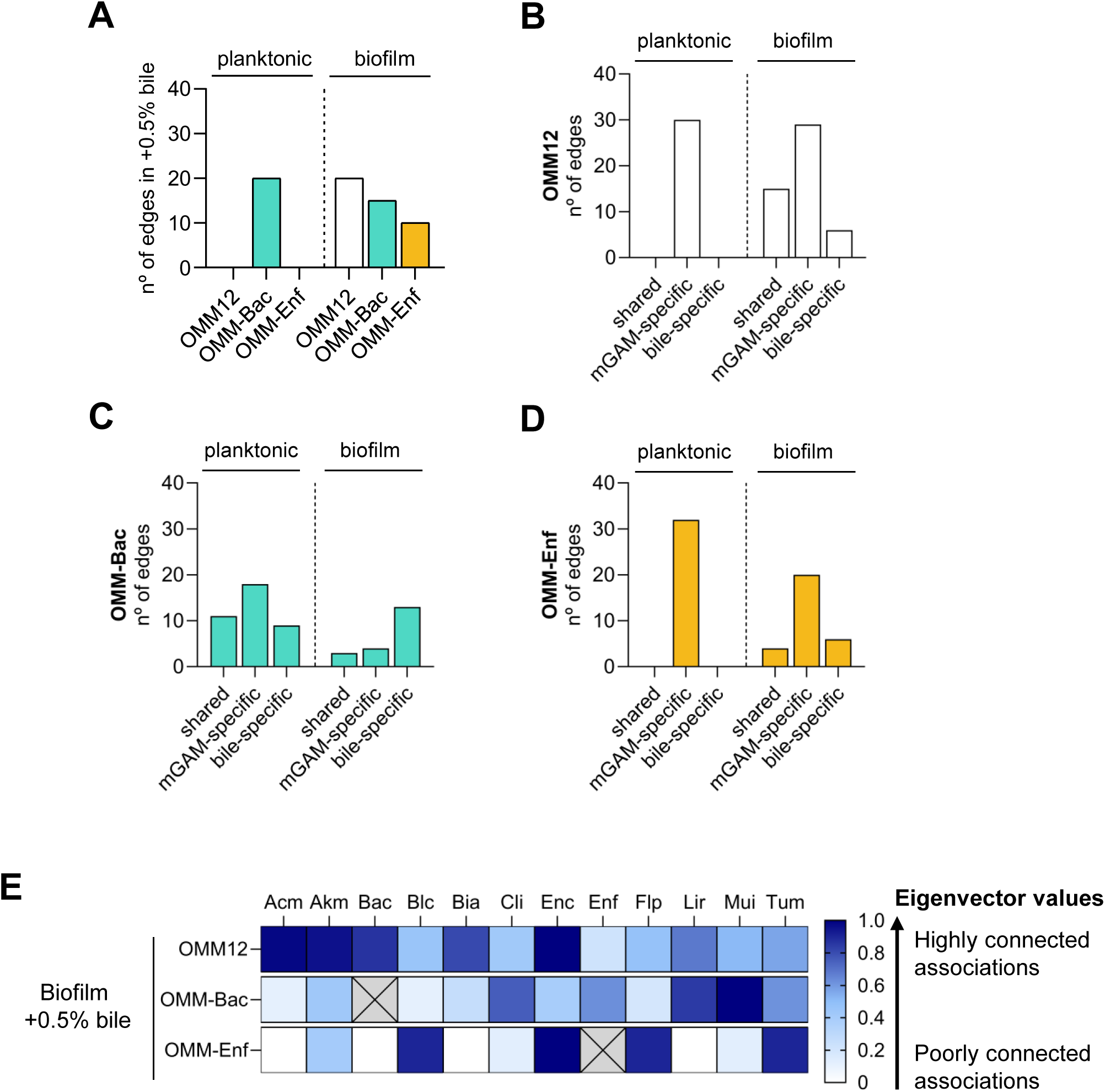
Network analysis of OMM12 and OMM11 communities shown in Fig.4 in +0.5% bile conditions. (**A**) Total number of edges in OMM12, OMM-Bac and OMM-Enf planktonic and biofilm communities. (**B**) Comparison of number of edge pairs shared between mGAM and bile in OMM12, (**C**) OMM-Bac and (**D**) OMM-Enf planktonic and biofilm communities. (**E**) Heatmap of eigenvector values that describe node centrality of each species of OMM12 and OMM11 biofilm communities in mGAM supplemented with 0.5% bile conditions.

**Fig. S11.**
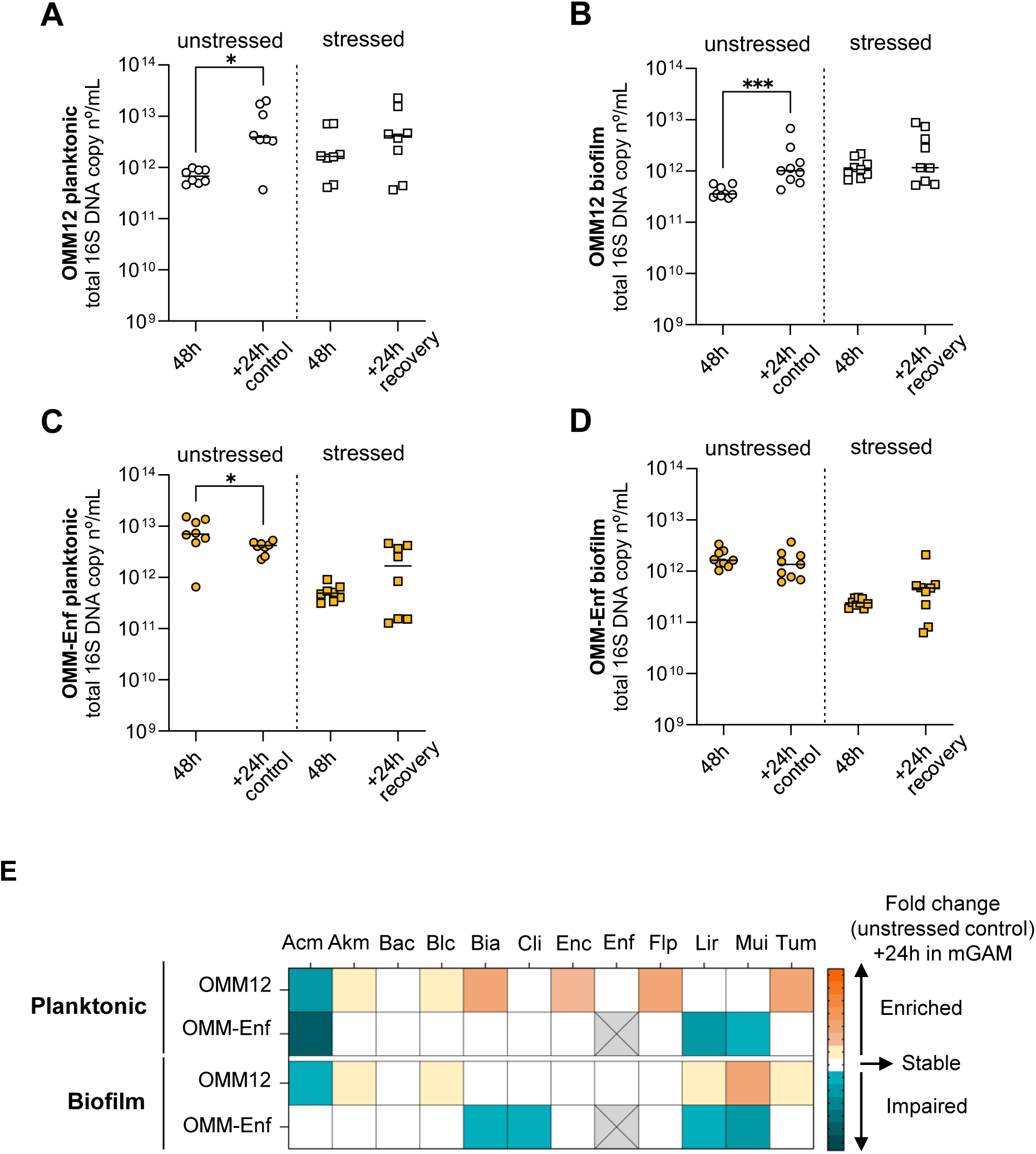
Total 16S DNA copy numbers and diversity index from OMM12 and OMM-Enf cultures from Fig5. (**A**) OMM12 planktonic and (**B**) biofilm cultures total 16S DNA copy numbers. (**C**) OMM-Enf planktonic and (**D**) biofilm cultures total 16S DNA copy numbers. (**E**) Heatmap representing the changes in abundance of each member after 24h exposed to fresh mGAM; fold change above 1 was enriched, between -1 and 1 was stable and below -1 was impaired. In A-D each dot represents the data from a culture replicate from 3 independent experiments. Data were analyzed with Mann-Whitney tests (* p<0.05; ** p<0.01; *** p<0.001).

**Fig. S12.**
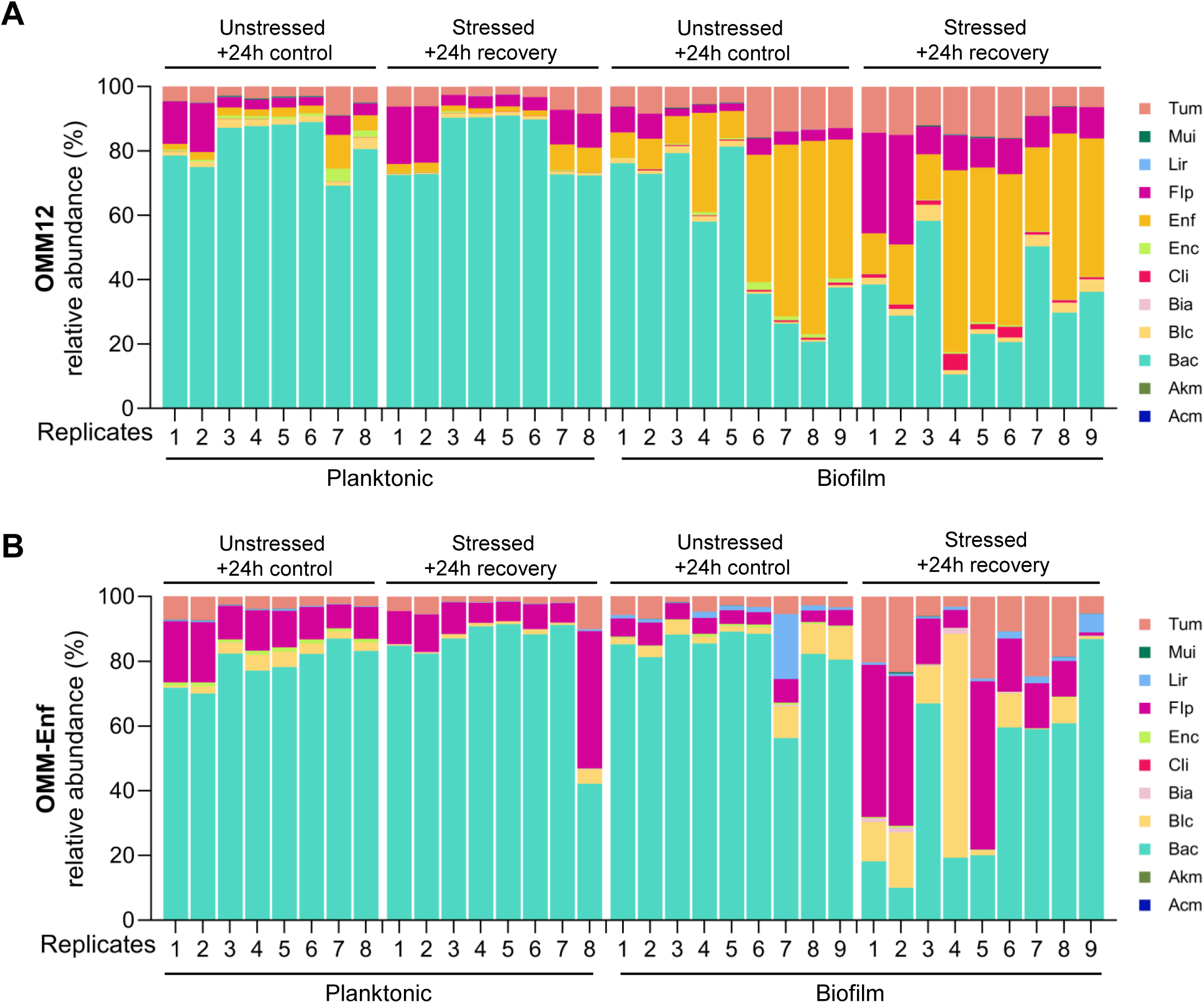
Community composition of OMM12 and OMM-Enf recovery phase cultures from Fig.5. (**A**) Relative abundance of OMM12 and (**B**) OMM-Enf communities. Bars show the data for each culture replicate.

**Fig. S13.**
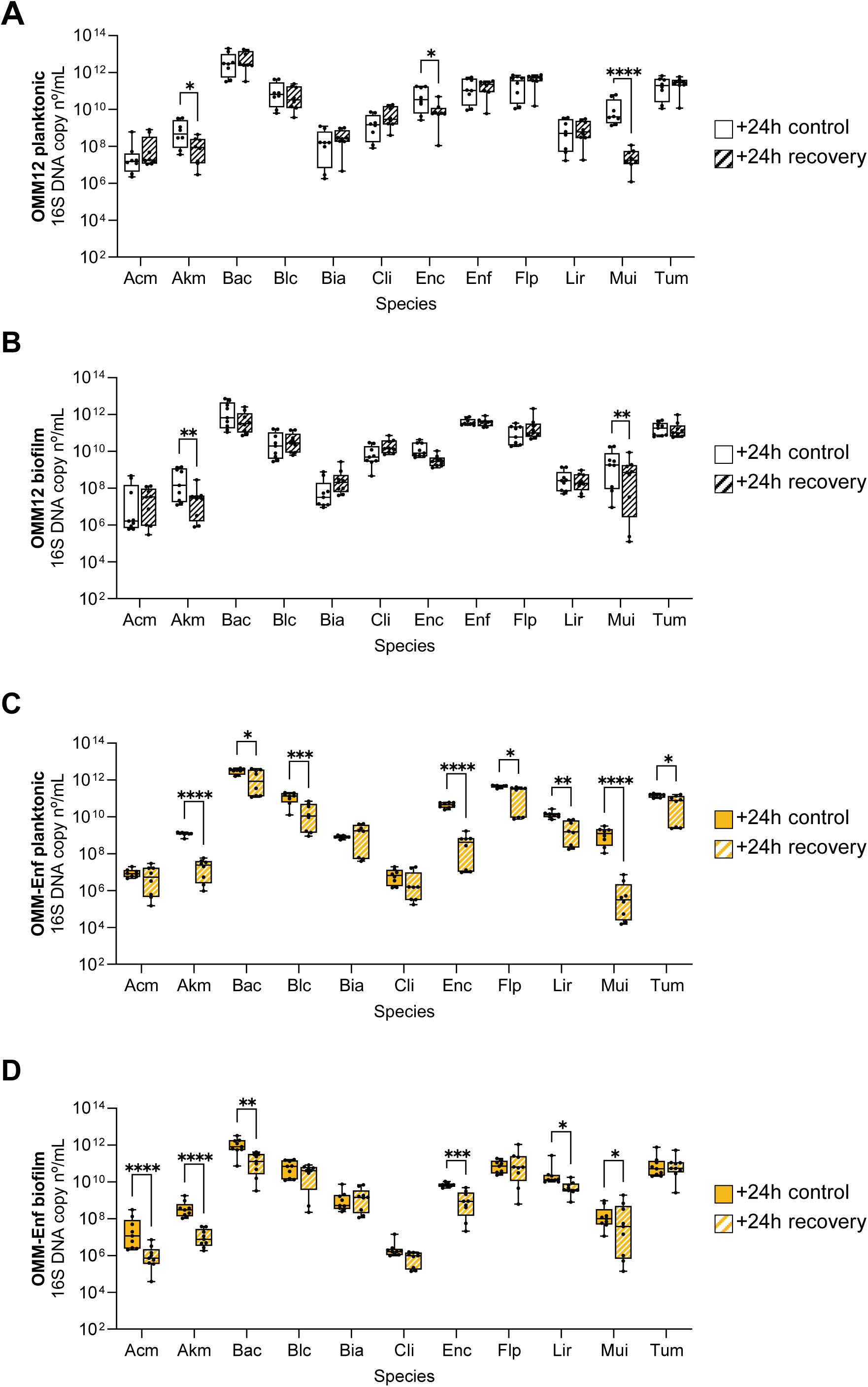
Absolute abundance of OMM12 and OMM-Enf communities shown in Fig.S12. (**A**) Absolute abundance of OMM12 planktonic and (**B**) biofilm communities. (**C**) Absolute abundance of OMM-Enf planktonic and (**D**) biofilm communities. Each dot represents the data from a culture replicate from 3 independent experiments. Comparison between each member’s abundance was analyzed with a linear mixed-effects model to determine statistical significance. Tukey adjustment was used to correct for multiple comparisons (* p<0.05; ** p<0.01; *** p<0.001; **** p<0.0001).

**Fig. S14.**
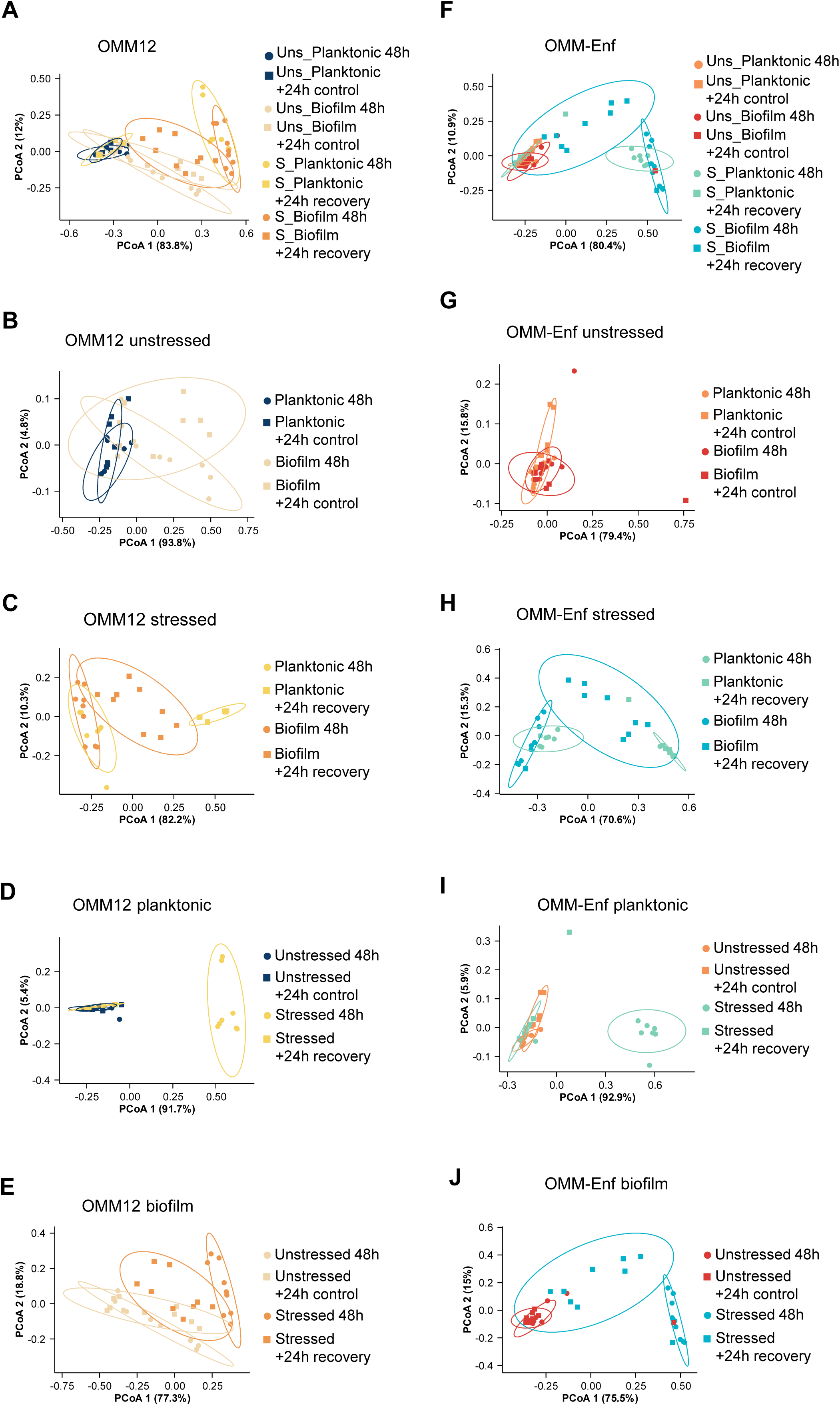
Community compositional shifts related to Fig.5. Principal Coordinate Analysis (PCoA) of Bray-Curtis dissimilarity index of (**A-E**) planktonic OMM12 and (**F-J**) OMM-Enf communities, ellipses with a 90% confidence interval for the first two coordinates of each group were drawn on the associated plot. Uns: unstressed, S: stressed. Each PCoA was tested with ANOSIM with Bonferroni correction; significance can be checked in the supplementary tables 1-10.

**Fig. S15.**
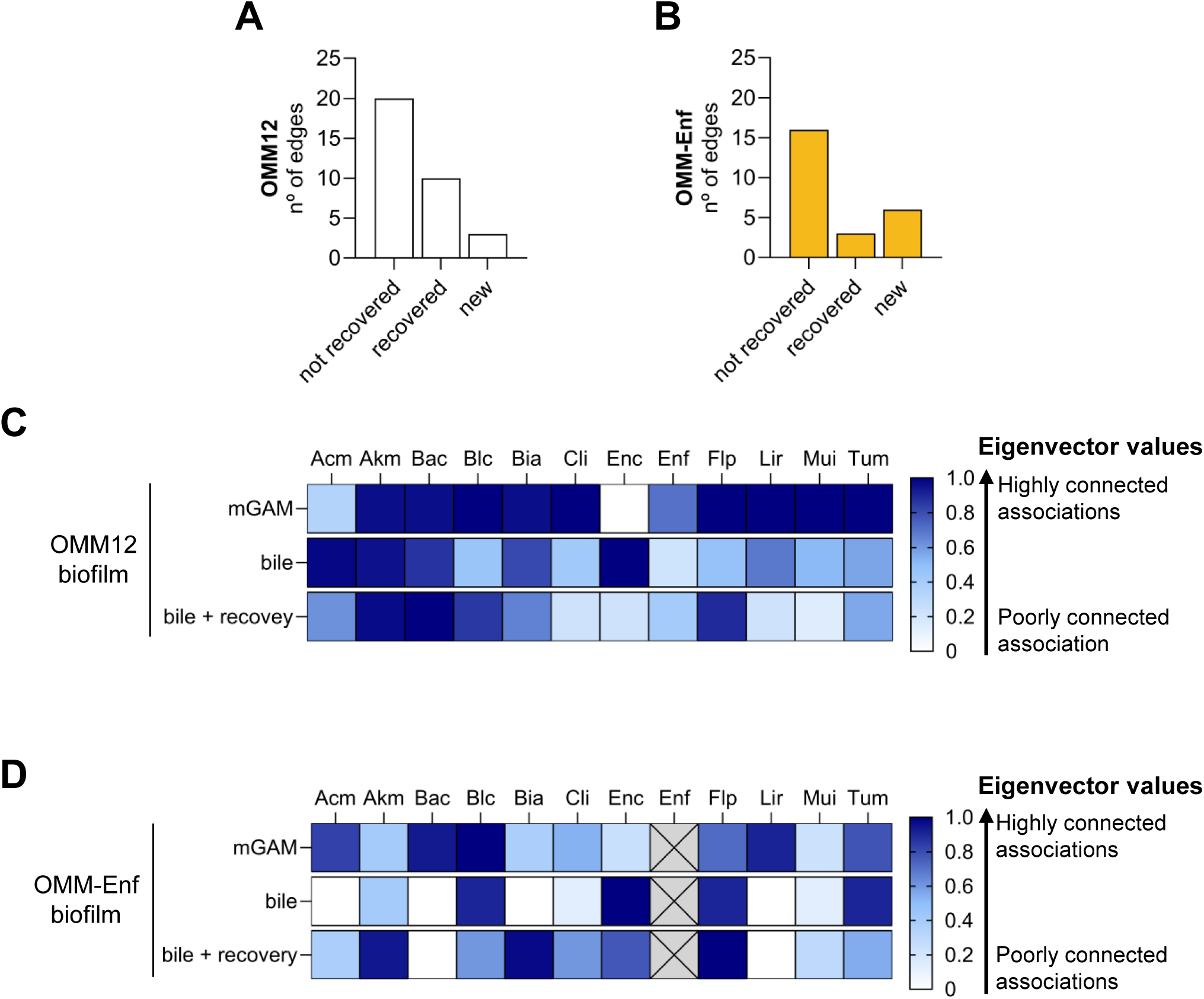
Network analysis of OMM12 and OMM-Enf recovered communities. (**A**) Comparison of number of edge pairs between bile + recovery and mGAM 48h in OMM12 and (**B**) OMM-Enf biofilm communities. Edge pairs were classified as: not recovered (present in mGAM, absent in bile, absent after stress removal); recovered (present in mGAM, absent in bile, reappeared after stress removal); or new (absent in both mGAM and bile, present only after stress removal). (**C**) Heatmap of eigenvector values that describe node centrality of each species of OMM12 biofilm communities in mGAM, upon bile exposure and after bile removal (bile + recovery). (**D**) Heatmap of eigenvector values that describe node centrality of each species of OMM-Enf biofilm communities in mGAM, upon bile exposure and after bile removal (bile + recovery). Data from mGAM 48h and bile correspond to the data shown in **Fig.S9D and Fig.S10E**, mGAM and +0.5% bile, respectively.

## REFERENCES

1. Buffie CG, Pamer EG. 2013. Microbiota-mediated colonization resistance against intestinal pathogens. Nat Rev Immunol 13:790–801.

2. Fan Y, Pedersen O. 2021. Gut microbiota in human metabolic health and disease. Nat Rev Microbiol 19:55–71.

3. Hooper LV, Littman DR, Macpherson AJ. 2012. Interactions Between the Microbiota and the Immune System. Science 336:1268–1273.

4. Turnbaugh PJ, Ley RE, Hamady M, Fraser-Liggett CM, Knight R, Gordon JI. 2007. The Human Microbiome Project. Nature 449:804–810.

5. Zeng Q, Feng X, Hu Y, Su S. 2025. The human gut microbiota is associated with host lifestyle: a comprehensive narrative review. Front Microbiol 16.

6. Bäckhed F, Ley RE, Sonnenburg JL, Peterson DA, Gordon JI. 2005. Host-Bacterial Mutualism in the Human Intestine. Science 307:1915–1920.

7. Eckburg PB, Bik EM, Bernstein CN, Purdom E, Dethlefsen L, Sargent M, Gill SR, Nelson KE, Relman DA. 2005. Diversity of the Human Intestinal Microbial Flora. Science 308:1635–1638.

8. Hou K, Wu Z-X, Chen X-Y, Wang J-Q, Zhang D, Xiao C, Zhu D, Koya JB, Wei L, Li J, Chen Z-S. 2022. Microbiota in health and diseases. Signal Transduct Target Ther 7:135.

9. Huttenhower C, Gevers D, Knight R, Abubucker S, Badger JH, Chinwalla AT, Creasy HH, Earl AM, FitzGerald MG, Fulton RS, Giglio MG, Hallsworth-Pepin K, Lobos EA, Madupu R, Magrini V, Martin JC, Mitreva M, Muzny DM, Sodergren EJ, Versalovic J, Wollam AM, Worley KC, Wortman JR, Young SK, Zeng Q, Aagaard KM, Abolude OO, Allen-Vercoe E, Alm EJ, Alvarado L, Andersen GL, Anderson S, Appelbaum E, Arachchi HM, Armitage G, Arze CA, Ayvaz T, Baker CC, Begg L, Belachew T, Bhonagiri V, Bihan M, Blaser MJ, Bloom T, Bonazzi V, Paul Brooks J, Buck GA, Buhay CJ, Busam DA, Campbell JL, Canon SR, Cantarel BL, Chain PSG, Chen I-MA, Chen L, Chhibba S, Chu K, Ciulla DM, Clemente JC, Clifton SW, Conlan S, Crabtree J, Cutting MA, Davidovics NJ, Davis CC, DeSantis TZ, Deal C, Delehaunty KD, Dewhirst FE, Deych E, Ding Y, Dooling DJ, Dugan SP, Michael Dunne W, Scott Durkin A, Edgar RC, Erlich RL, Farmer CN, Farrell RM, Faust K, Feldgarden M, Felix VM, Fisher S, Fodor AA, Forney LJ, Foster L, Di Francesco V, Friedman J, Friedrich DC, Fronick CC, Fulton LL, Gao H, Garcia N, Giannoukos G, Giblin C, Giovanni MY, Goldberg JM, Goll J, Gonzalez A, Griggs A, Gujja S, Kinder Haake S, Haas BJ, Hamilton HA, Harris EL, Hepburn TA, Herter B, Hoffmann DE, Holder ME, Howarth C, Huang KH, Huse SM, Izard J, Jansson JK, Jiang H, Jordan C, Joshi V, Katancik JA, Keitel WA, Kelley ST, Kells C, King NB, Knights D, Kong HH, Koren O, Koren S, Kota KC, Kovar CL, Kyrpides NC, La Rosa PS, Lee SL, Lemon KP, Lennon N, Lewis CM, Lewis L, Ley RE, Li K, Liolios K, Liu B, Liu Y, Lo C-C, Lozupone CA, Dwayne Lunsford R, Madden T, Mahurkar AA, Mannon PJ, Mardis ER, Markowitz VM, Mavromatis K, McCorrison JM, McDonald D, McEwen J, McGuire AL, McInnes P, Mehta T, Mihindukulasuriya KA, Miller JR, Minx PJ, Newsham I, Nusbaum C, O’Laughlin M, Orvis J, Pagani I, Palaniappan K, Patel SM, Pearson M, Peterson J, Podar M, Pohl C, Pollard KS, Pop M, Priest ME, Proctor LM, Qin X, Raes J, Ravel J, Reid JG, Rho M, Rhodes R, Riehle KP, Rivera MC, Rodriguez-Mueller B, Rogers Y-H, Ross MC, Russ C, Sanka RK, Sankar P, Fah Sathirapongsasuti J, Schloss JA, Schloss PD, Schmidt TM, Scholz M, Schriml L, Schubert AM, Segata N, Segre JA, Shannon WD, Sharp RR, Sharpton TJ, Shenoy N, Sheth NU, Simone GA, Singh I, Smillie CS, Sobel JD, Sommer DD, Spicer P, Sutton GG, Sykes SM, Tabbaa DG, Thiagarajan M, Tomlinson CM, Torralba M, Treangen TJ, Truty RM, Vishnivetskaya TA, Walker J, Wang L, Wang Z, Ward DV, Warren W, Watson MA, Wellington C, Wetterstrand KA, White JR, Wilczek-Boney K, Wu Y, Wylie KM, Wylie T, Yandava C, Ye L, Ye Y, Yooseph S, Youmans BP, Zhang L, Zhou Y, Zhu Y, Zoloth L, Zucker JD, Birren BW, Gibbs RA, Highlander SK, Methé BA, Nelson KE, Petrosino JF, Weinstock GM, Wilson RK, White O, The Human Microbiome Project Consortium. 2012. Structure, function and diversity of the healthy human microbiome. Nature 486:207–214.

10. Oliveira RA, Pamer EG. 2023. Assembling symbiotic bacterial species into live therapeutic consortia that reconstitute microbiome functions. Cell Host Microbe 31:472– 484.

11. Woelfel S, Silva MS, Stecher B. 2024. Intestinal colonization resistance in the context of environmental, host, and microbial determinants. Cell Host Microbe 32:820–836.

12. Bakkeren E, Piskovsky V, Foster KR. 2025. Metabolic ecology of microbiomes: Nutrient competition, host benefits, and community engineering. Cell Host Microbe 33:790–807.

13. Caballero-Flores G, Pickard JM, Núñez G. 2023. Microbiota-mediated colonization resistance: mechanisms and regulation. Nat Rev Microbiol 21:347–360.

14. Chavez-Arroyo A, Radlinski LC, Bäumler AJ. 2025. Principles of gut microbiota assembly. Trends Microbiol 33:718–726.

15. Culp EJ, Goodman AL. 2023. Cross-feeding in the gut microbiome: Ecology and mechanisms. Cell Host Microbe 31:485–499.

16. Wexler AG, Goodman AL. 2017. An insider’s perspective: Bacteroides as a window into the microbiome. Nat Microbiol 2:17026.

17. Burmølle M, Ren D, Bjarnsholt T, Sørensen SJ. 2014. Interactions in multispecies biofilms: do they actually matter? Trends Microbiol 22:84–91.

18. Motta J-P, Wallace JL, Buret AG, Deraison C, Vergnolle N. 2021. Gastrointestinal biofilms in health and disease. Nat Rev Gastroenterol Hepatol 18:314–334.

19. Béchon N, Ghigo J-M. 2022. Gut biofilms: Bacteroides as model symbionts to study biofilm formation by intestinal anaerobes. FEMS Microbiol Rev 46:fuab054.

20. Flemming H-C, Wuertz S. 2019. Bacteria and archaea on Earth and their abundance in biofilms. Nat Rev Microbiol 17:247–260.

21. Rumbaugh KP, Whiteley M. 2025. Towards improved biofilm models. Nat Rev Microbiol 23:57–66.

22. Ciofu O, Moser C, Jensen PØ, Høiby N. 2022. Tolerance and resistance of microbial biofilms. Nat Rev Microbiol 20:621–635.

23. Elias S, Banin E. 2012. Multi-species biofilms: living with friendly neighbors. FEMS Microbiol Rev 36:990–1004.

24. Flemming H-C, van Hullebusch ED, Neu TR, Nielsen PH, Seviour T, Stoodley P, Wingender J, Wuertz S. 2023. The biofilm matrix: multitasking in a shared space. Nat Rev Microbiol 21:70–86.

25. Edel M, Horn H, Gescher J. 2019. Biofilm systems as tools in biotechnological production. Appl Microbiol Biotechnol 103:5095–5103.

26. Philipp L-A, Bühler K, Ulber R, Gescher J. 2024. Beneficial applications of biofilms. Nat Rev Microbiol 22:276–290.

27. Duncan K, Carey-Ewend K, Vaishnava S. 2021. Spatial analysis of gut microbiome reveals a distinct ecological niche associated with the mucus layer. Gut Microbes 13:1874815.

28. Béchon N, Mihajlovic J, Vendrell-Fernández S, Chain F, Langella P, Beloin C, Ghigo J-M. 2020. Capsular Polysaccharide Cross-Regulation Modulates Bacteroides thetaiotaomicron Biofilm Formation. mBio 11:10.1128/mbio.00729-20.

29. Béchon N, Mihajlovic J, Lopes A-A, Vendrell-Fernández S, Deschamps J, Briandet R, Sismeiro O, Martin-Verstraete I, Dupuy B, Ghigo J-M. 2022. Bacteroides thetaiotaomicron uses a widespread extracellular DNase to promote bile-dependent biofilm formation. Proc Natl Acad Sci U S A 119:e2111228119.

30. Lopes A-A, Vendrell-Fernández S, Deschamps J, Georgeault S, Cokelaer T, Briandet R, Ghigo J-M. 2024. Bile-induced biofilm formation in Bacteroides thetaiotaomicron requires magnesium efflux by an RND pump. mBio 15:e03488–23.

31. Kister B, Viehof A, Rolle-Kampczyk U, Schwentker A, Treichel NS, Jennings SAV, Wirtz TH, Blank LM, Hornef MW, Bergen M von, Clavel T, Kuepfer L. 2023. A physiologically based model of bile acid metabolism in mice. iScience 26.

32. Peng, Wang S-H, Zhang Y-L, Chen M-Y, He K, Li Q, Huang W-H, Zhang W. 2024. Effects of bile acids on the growth, composition and metabolism of gut bacteria. Npj Biofilms Microbiomes 10:112.

33. Dubois T, Tremblay YDN, Hamiot A, Martin-Verstraete I, Deschamps J, Monot M, Briandet R, Dupuy B. 2019. A microbiota-generated bile salt induces biofilm formation in Clostridium difficile. Npj Biofilms Microbiomes 5:14.

34. Kelly SM, Lanigan N, O’Neill IJ, Bottacini F, Lugli GA, Viappiani A, Turroni F, Ventura M, van Sinderen D. 2020. Bifidobacterial biofilm formation is a multifactorial adaptive phenomenon in response to bile exposure. Sci Rep 10:11598.

35. Pumbwe L, Skilbeck CA, Nakano V, Avila-Campos MJ, Piazza RMF, Wexler HM. 2007. Bile salts enhance bacterial co-aggregation, bacterial-intestinal epithelial cell adhesion, biofilm formation and antimicrobial resistance of *Bacteroides fragilis*. Microb Pathog 43:78–87.

36. Zaidi AH, Bakkes PJ, Krom BP, van der Mei HC, Driessen AJM. 2011. Cholate-Stimulated Biofilm Formation by Lactococcus lactis Cells. Appl Environ Microbiol 77:2602–2610.

37. Olson JK, Navarro JB, Allen JM, McCulloh CJ, Mashburn-Warren L, Wang Y, Varaljay VA, Bailey MT, Goodman SD, Besner GE. 2018. An enhanced Lactobacillus reuteri biofilm formulation that increases protection against experimental necrotizing enterocolitis. Am J Physiol-Gastrointest Liver Physiol 315:G408–G419.

38. Sadiq FA, Wenwei L, Heyndrickx M, Flint S, Wei C, Jianxin Z, Zhang H. 2021. Synergistic interactions prevail in multispecies biofilms formed by the human gut microbiota on mucin. FEMS Microbiol Ecol 97:fiab096.

39. Wong JPH, Fischer-Stettler M, Zeeman SC, Battin TJ, Persat A. 2023. Fluid flow structures gut microbiota biofilm communities by distributing public goods. Proc Natl Acad Sci 120:e2217577120.

40. Brugiroux S, Beutler M, Pfann C, Garzetti D, Ruscheweyh H-J, Ring D, Diehl M, Herp S, Lötscher Y, Hussain S, Bunk B, Pukall R, Huson DH, Münch PC, McHardy AC, McCoy KD, Macpherson AJ, Loy A, Clavel T, Berry D, Stecher B. 2016. Genome-guided design of a defined mouse microbiota that confers colonization resistance against Salmonella enterica serovar Typhimurium. Nat Microbiol 2:16215.

41. Eberl C, Ring D, Münch PC, Beutler M, Basic M, Slack EC, Schwarzer M, Srutkova D, Lange A, Frick JS, Bleich A, Stecher B. 2020. Reproducible Colonization of Germ-Free Mice With the Oligo-Mouse-Microbiota in Different Animal Facilities. Front Microbiol 10.

42. Weiss AS, Niedermeier LS, von Strempel A, Burrichter AG, Ring D, Meng C, Kleigrewe K, Lincetto C, Hübner J, Stecher B. 2023. Nutritional and host environments determine community ecology and keystone species in a synthetic gut bacterial community. Nat Commun 14:4780.

43. Yilmaz B, Mooser C, Keller I, Li H, Zimmermann J, Bosshard L, Fuhrer T, Agüero MG de, Trigo NF, Tschanz-Lischer H, Limenitakis JP, Hardt W-D, McCoy KD, Stecher B, Excoffier L, Sauer U, Ganal-Vonarburg SC, Macpherson AJ. 2021. Long-term evolution and short-term adaptation of microbiota strains and sub-strains in mice. Cell Host Microbe 29:650–663.e9.

44. Osswald A, Wortmann E, Wylensek D, Kuhls S, Coleman OI, Peuker K, Strigli A, Ducarmon QR, Larralde M, Liang W, Treichel NS, Schumacher F, Volet C, Matysik S, Kleigrewe K, Gigl M, Rohn S, Guo C-J, Kleuser B, Liebisch G, Schnieke A, Ridlon JM, Bernier-Latmani R, Zeller G, Zeissig S, Haller D, Flisikowski K, Clavel T, Ocvirk S. 2025. Secondary bile acid production by gut bacteria promotes Western diet-associated colorectal cancer. Gut gutjnl-2024–332243.

45. Romero R, Zarzycka A, Preussner M, Fischer F, Hain T, Herrmann J-P, Roth K, Keber CU, Suryamohan K, Raifer H, Luu M, Leister H, Bertrams W, Klein M, Shams-Eldin H, Jacob R, Mollenkopf H-J, Rajalingam K, Visekruna A, Steinhoff U. 2022. Selected commensals educate the intestinal vascular and immune system for immunocompetence. Microbiome 10:158.

46. Kolland D, Kuhlmann M, de Almeida GP, Köhler A, Arifovic A, von Strempel A, Pourjam M, Bolsega S, Wurmser C, Steiger K, Basic M, Neuhaus K, Schmidt-Weber CB, Stecher B, Zehn D, Ohnmacht C. 2025. A specific microbial consortium enhances Th1 immunity, improves LCMV viral clearance but aggravates LCMV disease pathology in mice. Nat Commun 16:3902.

47. Weiss AS, Burrichter AG, Durai Raj AC, von Strempel A, Meng C, Kleigrewe K, Münch PC, Rössler L, Huber C, Eisenreich W, Jochum LM, Göing S, Jung K, Lincetto C, Hübner J, Marinos G, Zimmermann J, Kaleta C, Sanchez A, Stecher B. 2022. In vitro interaction network of a synthetic gut bacterial community. ISME J 16:1095–1109.

48. Derrien M, Vaughan EE, Plugge CM, De Vos WM. 2004. Akkermansia muciniphila gen. nov., sp. nov., a human intestinal mucin-degrading bacterium. Int J Syst Evol Microbiol 54:1469–1476.

49. Ronda C, Chen SP, Cabral V, Yaung SJ, Wang HH. 2019. Metagenomic engineering of the mammalian gut microbiome in situ. Nat Methods 16:167–170.

50. Hoyle BD, Costerton JW. 1991. Bacterial resistance to antibiotics: The role of biofilms, p. 91–105. In Salmon, JA, Garland, LG, Hoyle, BD, Costerton, JW, Seiler, N, Raeburn, D, Karlsson, J-A, Polak, A, Hartman, PG, Rohmer, M, Bisseret, P, Sutter, B, Burger, A, Jucker, E (eds.), Progress in Drug Research / Fortschritte der Arzneimittelforschung / Progrès des recherches pharmaceutiques. Birkhäuser, Basel.

51. Lee KWK, Periasamy S, Mukherjee M, Xie C, Kjelleberg S, Rice SA. 2014. Biofilm development and enhanced stress resistance of a model, mixed-species community biofilm. ISME J 8:894–907.

52. Pumbwe L, Skilbeck CA, Nakano V, Avila-Campos MJ, Piazza RMF, Wexler HM. 2007. Bile salts enhance bacterial co-aggregation, bacterial-intestinal epithelial cell adhesion, biofilm formation and antimicrobial resistance of Bacteroides fragilis. Microb Pathog 43:78–87.

53. Röttjers L, Faust K. 2018. From hairballs to hypotheses–biological insights from microbial networks. FEMS Microbiol Rev 42:761–780.

54. Garcia-Santamarina S, Kuhn M, Devendran S, Maier L, Driessen M, Mateus A, Mastrorilli E, Brochado AR, Savitski MM, Patil KR, Zimmermann M, Bork P, Typas A. 2024. Emergence of community behaviors in the gut microbiota upon drug treatment. Cell 187:6346–6357.e20.

55. Kumar M, Ji B, Zengler K, Nielsen J. 2019. Modelling approaches for studying the microbiome. Nat Microbiol 4:1253–1267.

56. Maier L, Goemans CV, Wirbel J, Kuhn M, Eberl C, Pruteanu M, Müller P, Garcia-Santamarina S, Cacace E, Zhang B, Gekeler C, Banerjee T, Anderson EE, Milanese A, Löber U, Forslund SK, Patil KR, Zimmermann M, Stecher B, Zeller G, Bork P, Typas A. 2021. Unravelling the collateral damage of antibiotics on gut bacteria. Nature 599:120– 124.

57. Dapa T, Ramiro RS, Pedro MF, Gordo I, Xavier KB. 2022. Diet leaves a genetic signature in a keystone member of the gut microbiota. Cell Host Microbe 30:183–199.e10.

58. Zheng X, Huang F, Zhao A, Lei S, Zhang Y, Xie G, Chen T, Qu C, Rajani C, Dong B, Li D, Jia W. 2017. Bile acid is a significant host factor shaping the gut microbiome of diet-induced obese mice. BMC Biol 15:120.

59. Stenman LK, Holma R, Korpela R. 2012. High-fat-induced intestinal permeability dysfunction associated with altered fecal bile acids. World J Gastroenterol 18:923–929.

60. Martens EC, Chiang HC, Gordon JI. 2008. Mucosal Glycan Foraging Enhances Fitness and Transmission of a Saccharolytic Human Gut Bacterial Symbiont. Cell Host Microbe 4:447–457.

61. Streidl T, Karkossa I, Segura Muñoz RR, Eberl C, Zaufel A, Plagge J, Schmaltz R, Schubert K, Basic M, Schneider KM, Afify M, Trautwein C, Tolba R, Stecher B, Doden HL, Ridlon JM, Ecker J, Moustafa T, von Bergen M, Ramer-Tait AE, Clavel T. 2021. The gut bacterium Extibacter muris produces secondary bile acids and influences liver physiology in gnotobiotic mice. Gut Microbes 13:1854008.

62. McMillan AS, Foley MH, Perkins CE, Theriot CM. 2023. Loss of Bacteroides thetaiotaomicron bile acid-altering enzymes impacts bacterial fitness and the global metabolic transcriptome. Microbiol Spectr 12:e03576–23.

63. Sannasiddappa TH, Lund PA, Clarke SR. 2017. In Vitro Antibacterial Activity of Unconjugated and Conjugated Bile Salts on Staphylococcus aureus. Front Microbiol 8.

64. Tian Y, Gui W, Koo I, Smith PB, Allman EL, Nichols RG, Rimal B, Cai J, Liu Q, Patterson AD. 2020. The microbiome modulating activity of bile acids. Gut Microbes 11:979–996.

65. O’Toole GA. 2011. Microtiter dish biofilm formation assay. J Vis Exp JoVE 2437.

